# Competition for the conserved branch point sequence influences physiological outcomes in pre-mRNA splicing

**DOI:** 10.1101/2024.10.09.617384

**Authors:** Karen Larissa Pereira de Castro, Jose M. Abril, Kuo-Chieh Liao, Haiping Hao, John Paul Donohue, William K. Russell, W. Samuel Fagg

**Affiliations:** Transplant Division, Department of Surgery, University of Texas Medical Branch, Galveston, TX, USA; RNA Genomics and Structure, Genome Institute of Singapore, Agency for Science, Technology, and Research (A*STAR) Singapore; Department of Biochemistry and Molecular Biology, University of Texas Medical Branch, Galveston, Texas, USA; Sinsheimer Labs, RNA Center for Molecular Biology, Department of Molecular, Cell and Developmental Biology, University of California, Santa Cruz, Santa Cruz, CA, USA

**Keywords:** alternative splicing, branch point, branch site, Quaking, RNA binding protein, splicing, Splicing Factor 1 (SF1)

## Abstract

Recognition of the intron branch point during spliceosome assembly is a multistep process that can influence mRNA structure and levels. A branch point sequence motif UACUAAC is variably conserved in eukaryotic genomes, but in some organisms, more than one protein can recognize it. Here, we show that SF1 and Quaking (QKI) compete for a subset of intron branch sites with the sequence ACUAA. SF1 activates exon inclusion through this sequence, but QKI represses the inclusion of alternatively spliced exons with this intron branch point sequence. Using mutant reporters derived from a natural intron with two branch site-like sequences, we find that when either branch point sequence is mutated, the other is utilized; however, when both are present, neither is used due to high-affinity binding and strong splicing repression by QKI. QKI occupancy at the dual branch site directly prevents SF1 binding and subsequent recruitment of spliceosome-associated factors. Finally, the ectopic expression of QKI in budding yeast (which lacks *QKI*) is lethal, at least in part due to the widespread repression of splicing. In conclusion, QKI can function as a splicing repressor by directly competing with SF1/BBP for a subset of branch point sequences that closely mirror its high-affinity binding site.

## Introduction

Up to 95% of human protein-coding genes can be alternatively spliced (1, 2). In contrast, only a minority of *S. cerevisiae* genes contain introns, and under normal growth conditions, most are efficiently spliced (3, 4). Many influences converge to promote extensive alternative splicing that is observed in higher eukaryotes, including genome complexity, *cis*-acting regulatory elements, and *trans*-acting factors (5–7). An early post-transcriptional step that is required for splicing is branch point (bp) recognition, which, along with 5′ splice site (ss), polypyrimidine (pY) tract, and 3′ss recognition, ultimately defines the spliceosome E-complex (8, 9). Sequence variability in these elements can influence the efficiency with which the E-complex and subsequent splicing complexes mature (10–15). An interesting bp sequence-specific observation is that different organisms maintain conservation of it; for example, in *Saccharomyces cerevisiae* (*S. cerevisiae*), it is nearly invariant (UACUAAC), while in mammals, it is more degenerate (YUNAY) (16–19). The underlying functional issues that explain this variability in bp sequence/branch site (bs) conservation are unknown. This suggests that bs variability might provide a way to control alternative splicing, but how this could be mediated is unclear.

RNA binding proteins (RBPs) can influence bs recognition by SF1 (mammals)(14, 20) or MSL5/BBP (*S. cerevisiae*)(14, 21). When these and other E-complex-associated proteins and snRNAs associate with the substrate, SF1/BBP is evicted and replaced by the SF3B complex, which recruits the 17S U2 snRNP, signaling maturation into the spliceosome A complex (22–24). Disruption of these events by either mutations or non-physiological concentrations of RBPs can lead to splicing defects and disease (25). For example, the balance between splicing-activating serine-arginine (SR) proteins versus splicing-repressing heterogeneous nuclear ribonucleoproteins (hnRNPs) regulates a large set of pre-mRNA substrates and provides an example of how competition between RBPs regulates alternative splicing (26, 27). However, it is unclear if RBP competition influences cell type-specific splicing or if it might have evolutionary implications.

The metazoan RBP <u>S</u>ignal <u>T</u>ransduction and <u>A</u>ctivator of <u>R</u>NA metabolism (STAR) family members possess a KH-type and QUA2 RNA binding domain and regulate disparate forms of RNA processing, including splicing (28). Interestingly, SF1 is a unique and the most divergent member of the STAR family as it lacks the QUA1 dimerization domain that all other members possess (20, 29, 30). The QUA1-containing members like Quaking (QKI), KHDRBS1-3, GLD1, and ASD2 exist as dimers and thus bind to a bipartite sequence motif in their target RNAs (31, 32). QKI (33, 34) and SF1 (20) are structurally similar (Fig 1A), and although SF1 lacks the QUA1 dimerization domain, they bind to a similar sequence motif. Interestingly, the SF1 binding motif (35) is more degenerate than the QKI consensus motif (31, 36–39), which is consistent with SF1’s role in the recognition of the degenerate mammalian bs. Analysis of tissue-specific splicing patterns nearly 20 years ago revealed that the UACUAAY motif is associated with muscle-specific exon skipping when located in the intron upstream of alternatively spliced exons or exon inclusion when located in the downstream intron, but the opposite pattern was observed in the brain (40). It was unclear how the “conserved bp sequence” UACUAAY might promote exon skipping or inclusion from the downstream intron. Still, it suggested that this element may have an additional and specialized role in tissue-specific alternative splicing. Clarity was provided to this model with the discovery that QKI directly regulates pre-mRNA splicing in C2C12 myoblasts via binding an ACUAAY element and exhibiting the prototypical “splicing code” positionally based regulatory pattern (38, 41, 42). This finding is also consistent with the observation that the nuclear isoform QKI5 is expressed at relatively high levels in muscle but at lower levels in the brain (38, 43–49). Thus, QKI5 enforces cell type-specific alternative splicing patterns by binding to a bp sequence that is also a high-affinity QKI binding site to promote exon skipping.

**Figure 1:**
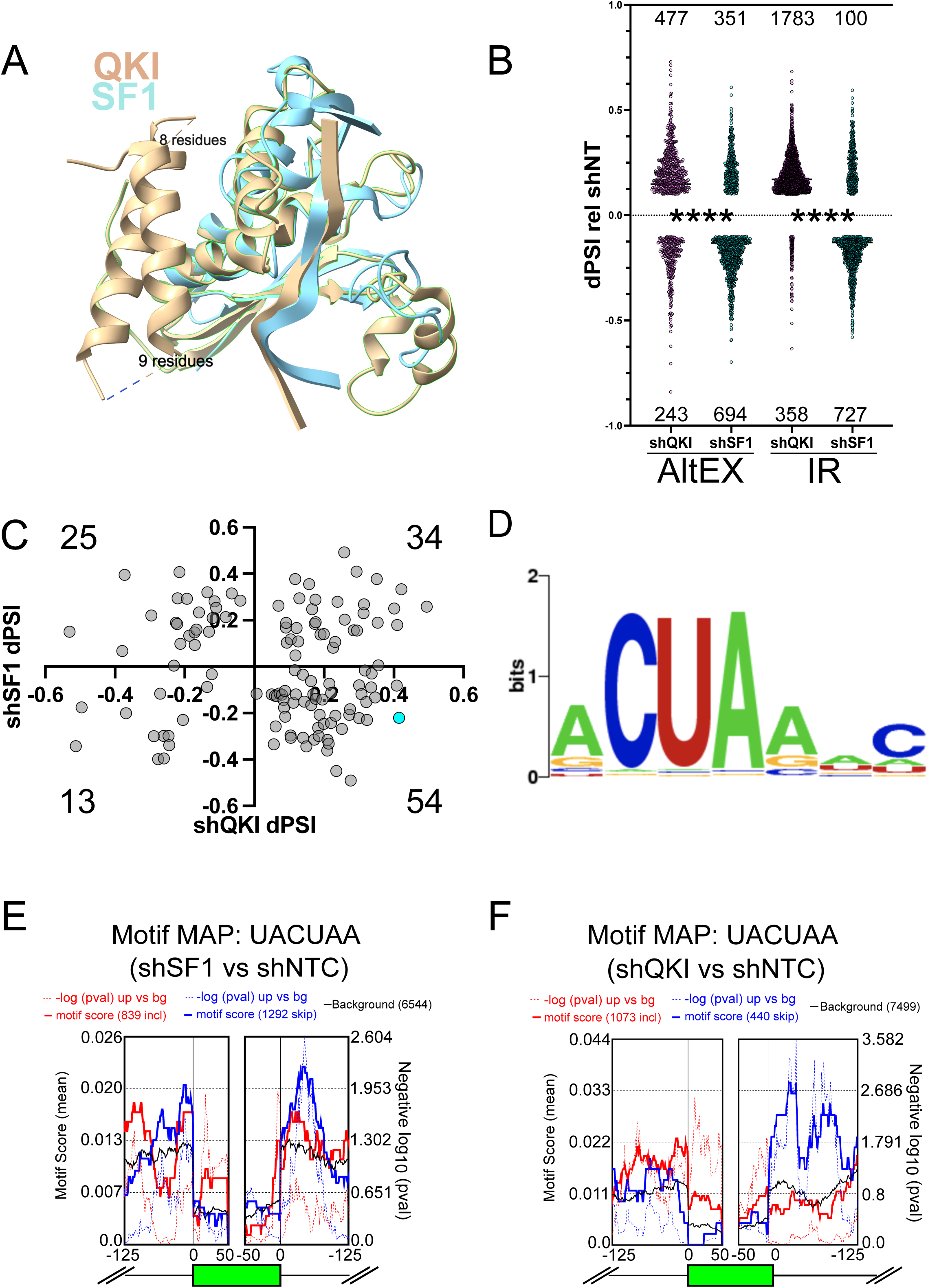
Similarity of SF1 and QKI and RNA-seq analysis of SF1 or QKI loss-of-function in HepG2 cells. A. Overlay of SF1 (cyan) and QKI (tan) KH and QUA2 protein domains: green highlighted region indicates KH and QUA2 domains of QKI; the two tan alpha helices on the left side make up the QUA1 domain in QKI (which SF1 lacks; text indicates residues in unstructured regions of QKI). B. Vast-tools analysis of the ENCODE RNA-seq data from SF1 (shSF1) or QKI shRNA (shQKI) knockdown compared to control shRNA (shNT) in HepG2 cells; the y-axis shows dPSI for shQKI or shSF1 relative to shNT for significantly altered alternatively spliced exons (AltEX; left) or intron retention (IR; right; dPSI ≥ |10| and MVdPSI > 0), and \*\*\*\**P* < 0.0001 by Mann-Whitney U when comparing the distribution of changes in shQKI relative to shNT to shSF1 relative to shNT). C. Scatter plot showing the distribution of AltEX events that changed under depletion of both SF1 (dPSI values relative to shNT on the y-axis) and QKI (dPSI values relative to shNT on the x-axis); the number in each quadrant indicates how many AltEX events were observed; *RAI14* exon 11 is shown in cyan. D. Simple Enrichment Analysis (SEA) of the intron region spanning 60 nt to 20 nt upstream of the 3’ss in QKI and SF1 regulated AltEX events shown in C (*P* < 0.05). E. rMAPS motif map for UACUAA generated for AltEX events changing during shSF1 compared to shNT by rMATS; motif scores (solid line) or -log10 P-value (dotted line) is shown in red for exons whose inclusion increases and in blue for exons whose inclusion decreases. F. rMAPS motif map as described in E. but for shQKI treatment compared to shNT.

Studies in lung/lung cells suggested that QKI could repress exon inclusion by competing with SF1 for a bs that might also function as a QKI binding site in the *NUMB* pre-mRNA (50). While this raised an intriguing possibility, it was unclear if these two RBPs were in direct competition for binding to a bona fide bs, and how competition for this specialized (but conserved consensus) bs might affect global splicing patterns, modulate splicing in naturally occurring intron substrates, alter spliceosome component recruitment, or influence evolutionary outcomes in splicing. Furthermore, SF1 knockdown in HeLa cells indicates it is required for viability and the splicing of several introns (51), but the extent to which its loss affects global splicing patterns is unknown. Here, we test these open questions and find that QKI5 can repress a set of alternatively spliced exons whose bs mirror the high-affinity QKI binding site. We use RAI14 exon 11 as a model and show that ACUAAC elements in intron 10 are true bs to which QKI or SF1 can bind to promote exon skipping or inclusion, respectively. Interestingly, QKI binding to RAI14 intron 10 bs leads to the recruitment of paraspeckle-associated proteins, while SF1 binding causes subsequent enrichment of the SF3a/b and U2 17S complexes. These discoveries expand the scope and clarify the bp competition model. Moreover, the addition of QKI5 into *S. cerevisiae* is lethal and concomitant with widespread splicing repression. Together, these findings suggest that the presence of *QKI* and degenerate bs may have co-evolved to expand the repertoire of cell type-specific alternative splicing in plants and multicellular eukaryotes.

## Results

### QKI and SF1 co-regulate a set of alternatively spliced exons through a distinct sequence motif

Based on analysis of their individual functions (14, 20, 35, 41, 52–56) and common structural features (Fig 1A), we hypothesized that QKI could repress the inclusion, while SF1 could activate the inclusion of a special subset of alternatively spliced exons that have ACUAA for their bs. To test this, we analyzed RNA sequencing (RNA-seq) datasets during QKI or SF1 knockdown (HepG2 cells treated with control non-targeting shRNA or shRNAs targeting *QKI* (shQKI) or *SF1* (shSF1)(57)) with VAST-tools to measure changes in alternative pre-mRNA splicing (58, 59). Consistent with potentially opposing functions in splicing, more exon inclusion and intron retention (IR) events are observed during QKI knockdown, but more exon skipping and fewer IR events are observed during SF1 knockdown; the distribution of these changes are significantly different when compared to one another (\*\*\*\**P* < 0.0001 by Mann-Whitney U; Fig 1B, and Supplemental Table S1; significance cutoff: change in percent spliced in (dPSI) > |10| and minimum value dPSI at 95% confidence interval (MVdPSI95) > 0). To measure the distribution of exons regulated by both QKI and SF1 (co-regulated exons), we plotted dPSI values of cassette exons under each condition relative to control for those that change under both knockdown conditions (dPSI > |10| in either shQKI or shSF1 relative to control, and MVdPSI95 > 0 in both shQKI relative to control and SF1 relative to control). Interestingly, we observed that 54 out of the 126 (43%) of the co-regulated alternatively spliced exons fell into the category where inclusion increases upon QKI knockdown and decreases upon SF1 knockdown (Fig 1C). Thus, the most highly represented set of these (by quadrant) is QKI-repressed and SF1-activated. We next asked if these co-regulated alternatively spliced exons are associated with any significantly enriched sequences in the upstream intron region in which the bs is found. To do so, we obtained the sequence from 63 to 20 nucleotides upstream of the 3′ splice site (these should include putative bs) but exclude pY tracts and 3′ splice sites) of these co-regulated exons or 1000 control exons from transcripts that are expressed (base mean > 100) but whose splicing is unchanged upon either QKI or SF1 knockdown (dPSI < |1|, MVdPSI = 0) and used Simple Enrichment Analysis (SEA (60)) to test if any motifs are significantly enriched. This analysis revealed the enrichment of a single ACUAA-like motif for the co-regulated exons (*P* = 0.033) but failed to identify any significantly enriched motifs from the control sequences (Fig 1D). Interestingly, this motif could potentially serve as a QKI (31, 36, 37) or SF1 (35, 57) binding site, or bs (17, 18). Finally, splicing analysis of these RNA-seq datasets using the complementary method rMATS (61) corroborated these findings by indicating more exon inclusion when QKI is knocked down and more skipping when SF1 is reduced (Fig 1E and 1F; Supplemental Tables S2 and S3, respectively). Motif enrichment analysis using rMAPS2 (62, 63) shows significant enrichment of the UACUAA motif in introns upstream of exons that are skipped more when SF1 is reduced compared to control (Fig 1E), and enrichment of this motif in introns upstream of exons more included when QKI is reduced compared to control (Fig 1F). In summary, the most common set of exons co-regulated by SF1 and QKI are SF1-activated and QKI-repressed, and these have enrichment of a bs-like sequence in their proximal upstream intron that also appears to be a QKI binding motif.

### RAI14 exon 11 is QKI-repressed and SF1-activated, with dual putative ACUAAC branch sites

To further investigate how competition between QKI and SF1 for the bs is mediated, we sought a prototypical alternatively spliced exon subject to this form of regulation. The criteria by which we narrowed our search were 1) a QKI-repressed and SF1-activated cassette exon with 2) experimental evidence of a functional ACUAAY bs and 3) experimental evidence of direct QKI binding. We previously found that QKI5 represses *Rai14* exon 11, and that reads from QKI iCLIP-seq map to the intron region just upstream of this exon in mouse myoblasts (38). Its inclusion is also repressed by QKI and activated by SF1 in HepG2 cells (Fig 1C (cyan dot) and Fig 2A; (57)). Inspection of the upstream intron region proximal to *RAI14* exon 11 revealed intriguing features that fulfilled the above criteria: tandem ACUAAC elements 34 nt or 43 nt upstream of the 3′ss, experimental evidence indicating that either of these could be used as a bs (18), and QKI eCLIP peaks from experiments using HepG2 or K562 cells (57, 64) that overlap with these putative ACUAAC bs sequences (Fig 2A). We next tested if the co-regulation of *RAI14* exon 11 splicing by QKI and SF1 observed in HepG2 cells could be observed in additional cell types. We measured *RAI14* exon 11 inclusion in HEK293 *QKI* KO cells and found a significant increase in its inclusion in the KO cells compared to WT (*P* < 0.0001 by Student’s t-test; Fig 2B). To reduce SF1 levels, we used two independent siRNAs targeting it in WT HEK293 cells. The first failed to produce an appreciable knockdown, but the second or the two combined reduced the SF1 protein level, which caused more *RAI14* exon 11 skipping (Fig 2C). Overexpression of myc:QKI5 but not a myc:QKI5 construct with reduced RNA binding activity (K120A;R124A (33)) promotes skipping of *RAI14* exon 11 in WT HEK293 cells, and overexpressing SF1 causes more inclusion (Fig S2). Although the physiological level of *Rai14* exon 11 inclusion is low in mouse myoblasts (perhaps due to the high level of QKI5 in these cells), knocking down *QKI5* or *SF1* in C2C12 cells leads to a significant increase or decrease, respectively, in its inclusion compared to control (*P* < 0.0001 by Student’s t-test; Fig 2D). Therefore, *RAI14* exon 11 splicing is repressed by QKI and activated by SF1 in various cell types, possibly by binding dual ACUAAC bs elements.

**Figure 2:**
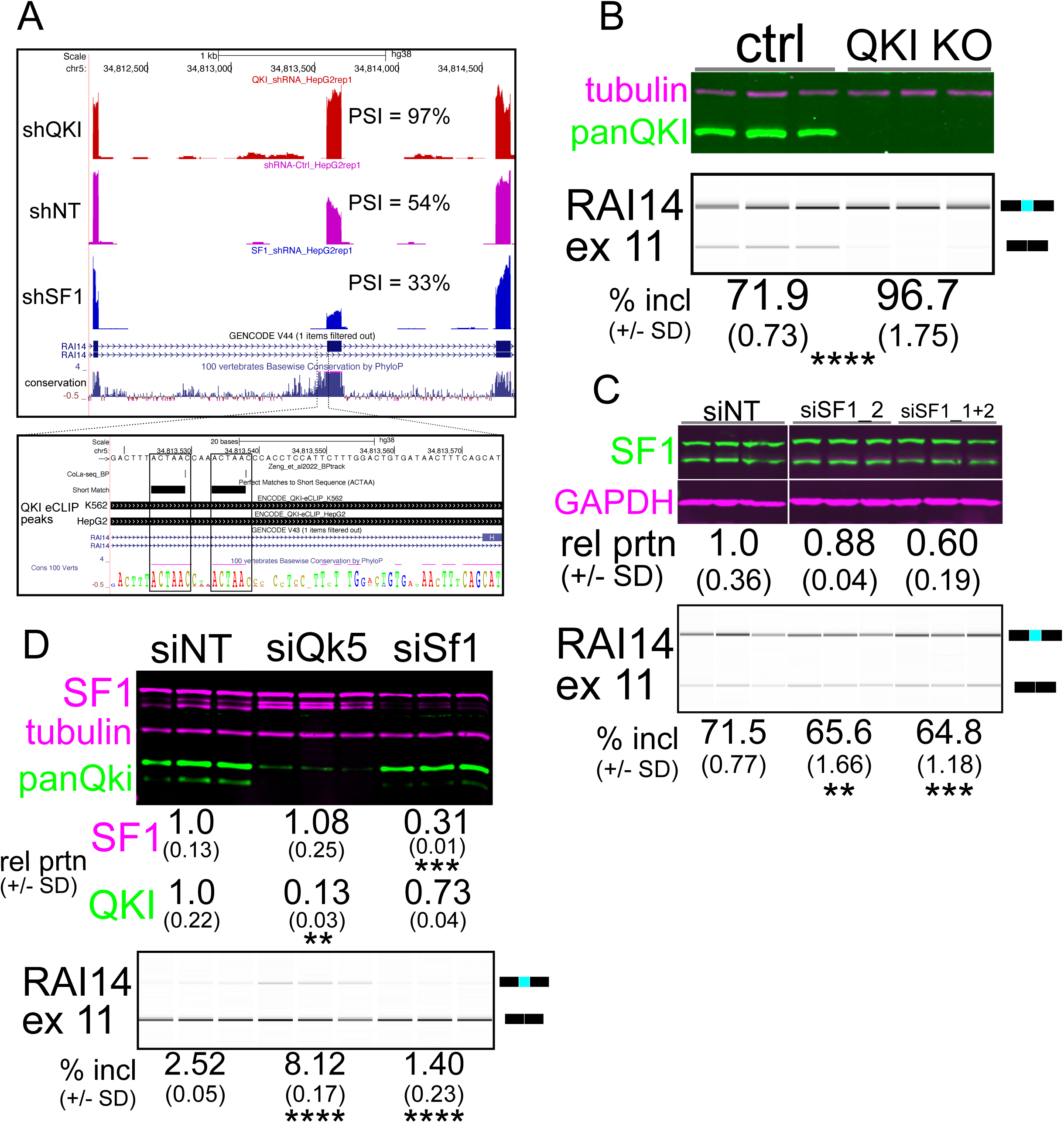
RAI14 exon 11 is repressed by QKI and activated by SF1. A. UCSC Genome Browser screenshot (top panel) with RNA-seq reads mapping to RAI14 for shQKI (top), shNT (middle), or shSF1 (bottom); percent spliced in (PSI) values are measured by Vast-tools; the inset shows boxed regions of (from top) two ACUAAC elements with Cola-seq branchpoints mapping to one nucleotide downstream of a branchpoint adenine, ACUAA elements by oligomatch, then QKI eCLIP peaks from K562 (top) or HepG2 (bottom) cells that overlap with these; conservation of 100 vertebrates is shown at the bottom. B. Western blot of protein extracted from HEK293 *QKI* KO cells (top) probed with anti-tubulin (magenta) or anti-panQKI (green) antibodies; RT-PCR of *RAI14* exon 11 and BioAnalyzer gel-like image (bottom) of RNA extracted from the cells above showing mean percent included ± standard deviation below (n = 3 biological replicates; \*\*\*\**P* < 0.001 by Student’s t-test compared to ctrl). C. Western blot (top) of proteins extracted from HEK293 cells transfected with siNT, siSF1_2 or siSF1_1+2, probed with anti-SF1 (green) or anti-Gapdh (magenta) antibodies; the protein abundance (fold change relative to the siNT control ± standard deviation) is shown below; RT-PCR of *RAI14* exon 11 and BioAnalyzer gel-like image (bottom) of RNA extracted from the cells described above with mean percent included ± standard deviation below (n = 3 biological replicates; \*\**P* < 0.01 or \*\*\**P* < 0.001 by Student’s t-test compared to siNT). D. Western blot (top) of proteins extracted from C2C12 myoblasts transfected with siNT, siQki or siSf1_1+2, probed with anti-SF1 (magenta; top), anti-tubulin (magenta; middle) or anti-panQki (green; bottom). The protein abundance (fold change relative to the siNT control ± standard deviation is shown below (n = 3 biological replicates; \*\**P* < 0.01 or \*\*\**P* < 0.001 by Student’s t-test); RT-PCR of *Rai14* exon 11 and BioAnalyzer gel-like image (bottom) of RNA extracted from the C2C12 cells described above with mean percent included ± standard deviation indicated below (\*\*\*\**P* < 0.0001 by Student’s t-test).

### Efficient RAI14 exon 11 skipping requires tandem ACUAAC elements, either of which can be used as a bp

Next, we asked how the ACUAAC elements in *RAI14* intron 10 influence exon 11 alternative splicing. Previous investigation did not stringently discriminate between whether QKI and SF1 competed directly for binding to the bs or if dimeric QKI bound to a bipartite motif that flanked the bs and thus occluded SF1 binding to the true bp (50). We hypothesized that: 1) efficient exon 11 skipping/splicing repression requires two intact ACUAAC elements (or at least a single ACUAAC element and QKI “half-site” (which would constitute the previously SELEX-defined high affinity Quaking response element (QRE)) (31)), 2) one ACUAAC element or the other must be present for any exon inclusion (either can be a bs), 3) loss of both ACUAAC elements would lead to no exon inclusion (one or the other is required to have a bs), 4) conversion of ACUAAC elements to another (non-QKI binding motif) bs would lead to more inclusion; regarding RNA stability: 5) removal of either ACUAAC element could destabilize the transcript, but 6) restoring either to a UAAC “half-site” could rescue this defect and promote exon skipping. To test these, we generated a splicing reporter by cloning 243 nucleotides of RAI14 intron 10, RAI14 exon 11, and 95 nucleotides of RAI14 intron 11 into the beta globing splicing reporter pDUP51 (DUP-RAI14 exon 11). We first deleted either the first, the second, or both ACUAAC elements (Fig 3A), transfected these reporter plasmids into C2C12 cells, and then measured exon inclusion by RT-PCR or total transcript stability by RT-qPCR. Deletion of either the upstream or downstream ACUAAC element causes ∼40% increase in inclusion (Fig 3B), indicating that both sites are required for strong splicing repression. Interestingly, we the appearance of a mis-spliced product from the upstream deletion mutant compared to WT (Fig 3B); the latter may be due to the use of a weak/cryptic bp and 3′ss (see * on Fig 3B). Essentially no inclusion of RAI14 exon 11 is observed in the absence of both ACUAAC elements, except for the mis-spliced product noted above (Fig 3B). Control experiments show that these amplicons are reverse transcriptase-dependent, indicating that the products detected are not due to plasmid DNA or PCR product contamination, and corroborate the results described above (Suppl Fig S3). Moreover, loss of the upstream ACUAAC element causes markedly reduced total reporter RNA levels; we observe a similar trend in the other deletion mutants, but to a lesser magnitude (Fig 3C). Therefore, potent splicing repression of RAI14 exon 11 requires both ACUAAC elements, and one or the other is required for any exon inclusion, and thus they are bona fide bs.

**Figure 3:**
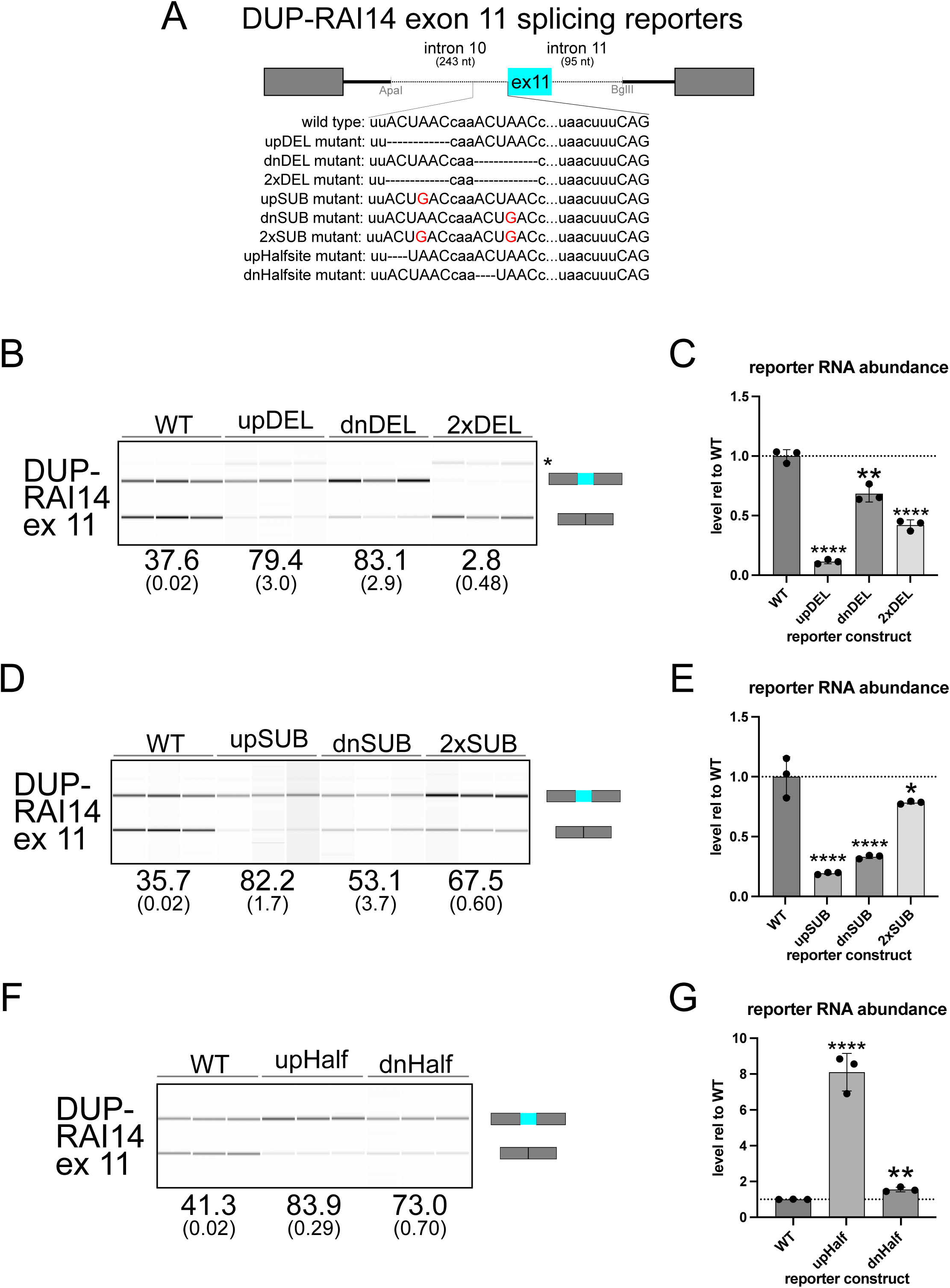
Analysis of DUP-RAI14 exon 11 (ex 11) splicing reporter. A. Schematic of the beta globin pDUP-RAI14 ex 11 splicing reporters indicating the region of intron 10, exon 11, and intron 11 included; the inset below show the different constructs (wild type or mutant) tested. B. RT-PCR and BioAnalyzer gel-like image from RNA extracted from C2C12 cells transfected with *RAI14* ex 11 wild type or deletion mutant reporters; mean percent included values are shown below (+/- the standard deviation; *indicates an unidentified/spurious product). C. RT-qPCR measuring total reporter RNA level, normalized to *Eef1a1*, from RNA described in B., and shown as fold-change relative to WT (\*\**P* < 0.01 or \*\*\*\**P* < 0.001 by Student’s t-test). D. RT-PCR and BioAnalyzer gel-like image from RNA extracted from C2C12 cells transfected with *RAI14* ex 11 wild type or substitution mutant reporters, analyzed as described in B. E. RT-qPCR measuring total reporter RNA level, normalized to *Eef1a1*, from RNA described in D., and shown as fold-change relative to WT (\**P* < 0.05 or \*\*\*\**P* < 0.001 by Student’s t-test). F. RT-PCR and BioAnalyzer gel-like image from RNA extracted from C2C12 cells transfected with *RAI14* ex 11 wild type or half-site mutant reporters, analyzed as described in B (\*\*\**P* < 0.001, \*\*\*\**P* < 0.0001). G. RT-qPCR measuring total reporter RNA level, normalized to *Eef1a1*, from RNA described in F., and shown as fold-change relative to WT (\*\**P* < 0.01 or \*\*\*\**P* < 0.001 by Student’s t-test). Each experiment was conducted in biological triplicate, and for RT-PCR with BioAnalyzer measurement, each comparison compared to WT reporter showed *P* < 0.0001 by Student’s t-test.

We next asked whether substituting either ACUAAC element, or both, to ACU*G*AC (which should be a poor substrate for QKI binding but a suitable bp sequence (Fig 3A)) would also lead to increased inclusion of RAI14 exon 11 or influence reporter RNA stability. We observe a large increase in exon inclusion and decrease in skipping for either of the single substitution mutants and for the double substitution mutant (∼20-35%; Fig 3D). As above, the amplification of these products is RT-dependent (Suppl Fig S3). Transcript abundance is significantly reduced in either single substitution mutant (\*\*\*\**P* < 0.0001 by Student’s t-test), and is slightly lower in the double substitution mutant (Fig 3E). Finally, converting either ACUAAC element to a UAAC “half-site” results in comparable increases in inclusion/decreases in skipping compared to WT as described in the deletion or substitution mutants (Fig 3F), which are also RT-dependent (Suppl Fig S3). Interestingly, these “half-site” mutants show a higer level of abundance than the WT reporter, especially in the upstream “half-site” mutant (Fig 3G). In summary, removal of either ACUAAC element or conversion to ACUGAC leads to splicing activation of RAI14 exon 11, indicating a requirement for dual ACUAAC elements for potent splicing repression.

### QKI binding to Rai14 intron 10 requires both ACUAAC elements and prevents spliceosome recruitment

We next hypothesized that QKI binding to Rai14 intron 10 requires both ACUAAC elements, that its binding would prevent spliceosome recruitment by blocking SF1, and that removal of either ACUAAC element would favor SF1 binding and recruitment of spliceosome components. To minimize bias, we initially performed RNA affinity chromatography (RAC) using 64 nt of intron sequence upstream of the 3′ss and including 6 nt of exonic sequence linked to a tobramycin aptamer (WT)(65), and the same sequence but with either the upstream ACUAAC (upDEL), downstream ACUAAC (dnDEL), or both ACUAACs (2xDEL), as well as a tobramycin aptamer (APT) only RNA (Fig 4A). Our rationale for this was to identify all proteins bound to these substrates and correlate them with splicing outcomes: WT shows low levels of inclusion, upDEL slightly higher levels of inclusion, dnDEL high levels of inclusion, and 2xDEL lacks a bp, and so is completely skipped. We added C2C12 myoblast nuclear extract (NE) to these under conditions that would favor splicing (with ATP) and then collected the associated proteins and identified them by liquid chromatography with tandem mass spectrometry (LC-MS/MS) (65). Quaking is readily identified as associating with the WT RAC substrate and in the input NE but is undetectable in each of the mutant RAC substrates and the APT-only control; SF1 is not detected associating with any RAC substrate but was present in NE (Fig 4B and 4C, and Supplemental Table S4). We used previously published LC-MS/MS datasets to generate a list of early spliceosome (E complex) and 17S U2 snRNP components (E/U2) (66–68) in order to focus on these proteins in our RAC-LC-MS/MS datasets to test our hypotheses. Subsequently, we found that the proteomic profiles detected in association with the WT and 2xDEL substrates are the most similar (Fig 4B), suggesting that QKI protein binding or loss of bs leads to similar E/U2 protein binding patterns. Indeed, in either of the single deletion mutants, we observe more enrichment of E/U2 components with the substrate RNA (24 positive and 5 negative values in upDEL and 16 positive values and 7 negative values in dnDEL) while the WT and 2xDEL substrates show a more balanced distribution (18 positive and 12 negative or 18 positive and 11 negative values, respectively; Fig 4C). Western blot analysis using the same RAC approach also shows robust QKI association with the WT substate, and undetectable or nearly undetectable SF1 protein, validating our LC-MS/MS findings (Fig 4D).

**Figure 4:**
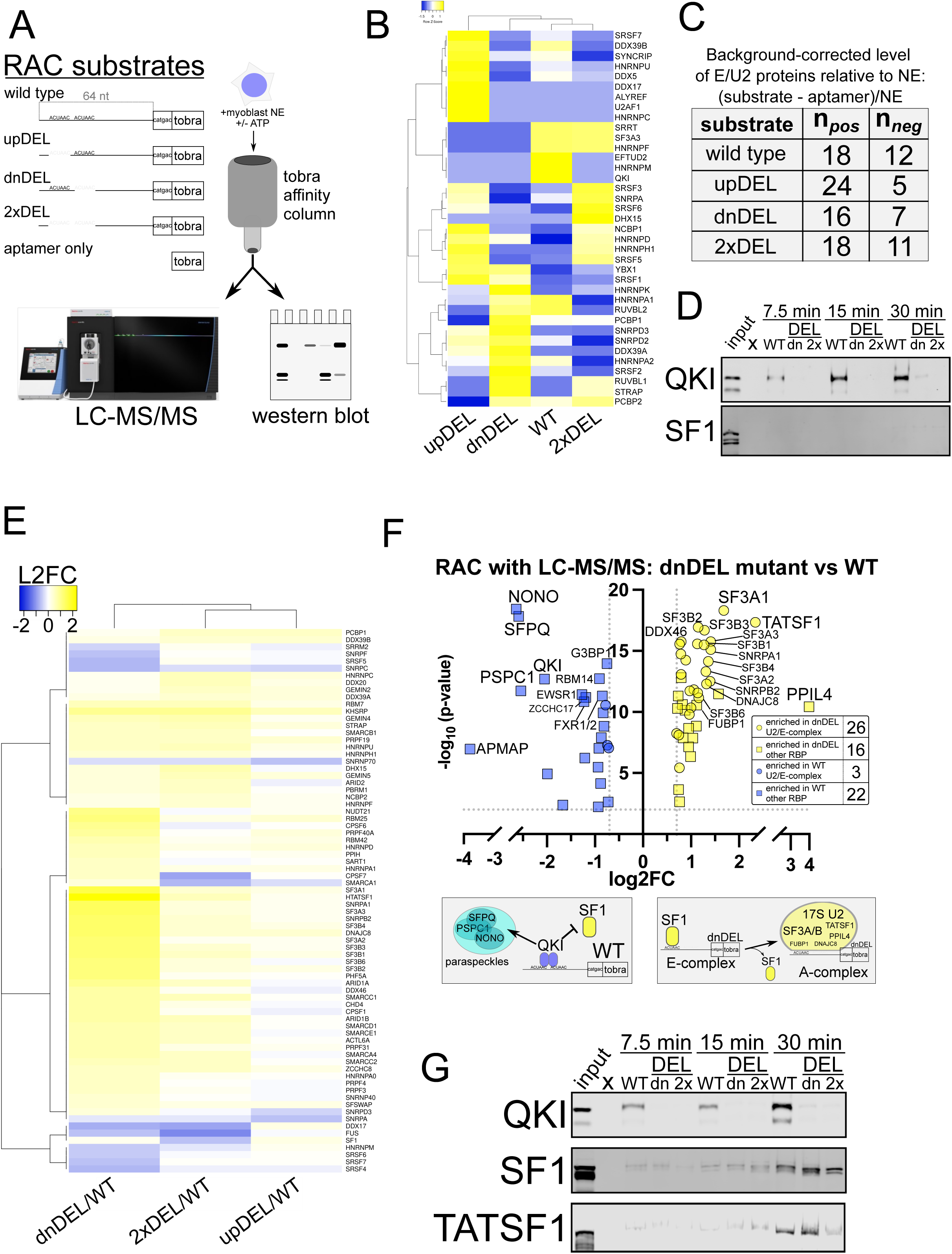
Analysis of the proteins associating with *RAI14* intron 10 RNA. RNA affinity chromatography (RAC) with liquid chromatography and tandem mass spec or western blot analysis. A. Schematic representation of substrates used for RNA affinity chromatography (RAC) which included 64 nt of *RAI14* intron 10, 6 nt of exon sequence, and the tobramycin aptamer (tobra): wild type, upDEL, dnDEL, 2xDEL and aptamer only; C2C12 nuclear extract (NE) was incubated with these, and RAC-associated eluates were analyzed by liquid chromatography with tandem mass spectrometry (LC-MS/MS) or western blot. B. Heatmap showing hierarchical clustering of early spliceosome and 17S U2 snRNP protein (E/U2) abundance detected in the RAC-LC-MS/MS datasets for each substate shown and in the presence of ATP; these represent background-corrected levels relative to NE (see Methods) and the scale bar shows row Z-score values. C. Numbers observed for relative levels of E/U2 proteins (background corrected relative to NE) observed associating with each RAC substate with either a positive (n_pos_) or negative (n_neg_) value. D. Western blot of NE (input) or WT, dnDEL (dn), or 2xDEL (2x) RAC time-course (+ATP as described in B and C) for 7.5 minutes (left), 15 minutes (middle), or 30 minutes (right) probed with anti-panQki (top) or anti-SF1(bottom) antibodies. E. Heatmap showing hierarchical clustering of early spliceosome and 17S U2 snRNP protein (E/U2) abundance detected in the RAC-LC-MS/MS datasets for each substate shown and in the absence of ATP; these represent data-independent acquisition (DIA; see methods) values normalized to NE and each mutant is shown as log2 fold change relative to the WT substate and passed cutoff of log_2_ fold change > |0.2| and *P* < 0.01. F. Volcano plot comparing LC-MS/MS log_2_ protein abundance (log_2_ fold-change (log2FC); x-axis) of E/U2 (circles) and other RBPs (squares) observed associating with RAC substates dnDEL compared to WT (y-axis shows -log_10_ *P*-value) of enriched proteins (cutoff: L2FC > |0.7| and *P* < 0.01); inset shows the number observed for those enriched in dnDEL (yellow) or WT (blue); schematic below shows model of RAC substates recruitment to distinct protein-associated species. G. Western blot of NE (input) or WT, dnDEL (dn), or 2xDEL (2x) RAC time-course (-ATP as described in E and F) for 7.5 minutes (left), 15 minutes (middle), or 30 minutes (right) probed with anti-panQki (top), anti-SF1 (middle), or anti-TATSF1 (bottom).

Previous studies indicate that excluding ATP from RNA splicing/binding assays favors a more stable association of early splicing complexes, including SF1 (12, 69, 70). Therefore, we performed RAC-LC-MS/MS with the same substrates in the absence of ATP (Supplemental Table S5). We also employed a data-independent acquisition (DIA) LC-MS/MS method, utilizing NE to construct a peptide search library, thereby enhancing the specificity and sensitivity of detection (see Methods). We observe significant enrichment (log_2_ fold change > |0.2| and *P* < 0.01) of E/U2 components (including SF1) binding to the RNA in the absence of the downstream ACUAAC (dnDEL) element compared to WT RNA (Fig 4E). This trend is also observed but to a lesser extent in the absence of the upstream ACUAAC (upDEL) and in the absence of both (2xDEL) mutants relative to WT; the proteome profiles associated to these two substrates are also more similar to one another than the dnDEL comparison to WT, consistent with lower splicing efficiency or no splicing, respectively (Fig 4E). Closer and more stringent (log_2_ fold change ≥ |0.7| and *P* < 0.01) inspection of the proteins binding preferentially to the dnDEL mutant RNA compared to the WT RNA reveal enrichment of 26 E/U2 components along with 16 other annotated (non-E/U2) RBPs; we observe depletion (or enrichment in WT) of only 3 E/U2 proteins but of 22 other annotated RBPs including QKI (Fig 4F). Interestingly, some of the most enriched protein components associating with the dnDEL RNA are TATSF1, Sf3A1, PPIL4, Sf3A3, Sf3B4, DNAJC8, Sf3A2, Sf3B3, Sf3B1, and FUBP1 (Fig 4F) many of which are known to interact with the bs and SF1 or are members of the Sf3 complex, which promotes spliceosome maturation by displacing SF1 from the bp and then recruiting the U2 snRNP (24, 71–73). The most significantly enriched proteins associating with the WT RAC substate are NONO, SFPQ, PSPC1, and QKI; the former three are found in paraspeckles that are formed by liquid-liquid phase separation (74), and QKI has recently been identified as a paraspeckle component protein (75). Western blot analysis validates the above findings indicating that QKI associates with high affinity to the WT but not dnDEL or 2xDEL RNA, and, in the absence of ATP, SF1 and TATSF1 associate to a greater degree with the dnDEL mutant than either of the other RAC substates (Fig 4G). Therefore, both ACUAAC elements are required for QKI binding and repression of Rai14 exon 11 splicing; removing the downstream ACUAAC element causes SF1 binding, and the subsequent recruitment of the spliceosome A complex machinery and de-repression of RAI14 exon 11 splicing.

### Ectopic expression of QKI5 in S. cerevisiae is lethal and causes pre-mRNA splicing defects

Our results indicate that QKI and SF1 can directly compete for a subset of ACUAA bp sequences, so we hypothesized that expressing the QKI5 isoform in *S. cerevisiae* (where the bp sequence is nearly invariant UACUAAC and lacks *QKI*) would be lethal and cause defective splicing. To test this, we inserted a galactose-inducible cDNA encoding either EGFP, mutant K120A;R124A QKI5, or wildtype QKI5 into BY4741 at the *URA3* locus, and each grew normally on glucose-containing media where the transgene is repressed. Galactose-induction is tolerated for the EGFP- and mutant QKI5-containing yeast, but is lethal for the wildtype QKI5-containing strain (Fig 5A). Growth curve analysis indicates that after about 4h the QKI5-expressing BY4741 cells cease proliferation (Supplemental Fig S5A). Interestingly, by 24h most of these cells have a “large-budded” morphology, which suggests a defect in cell division; EGFP- and mutant QKI5-expressing cells grow normally (Supplemental Fig S5A and S5B).

**Figure 5:**
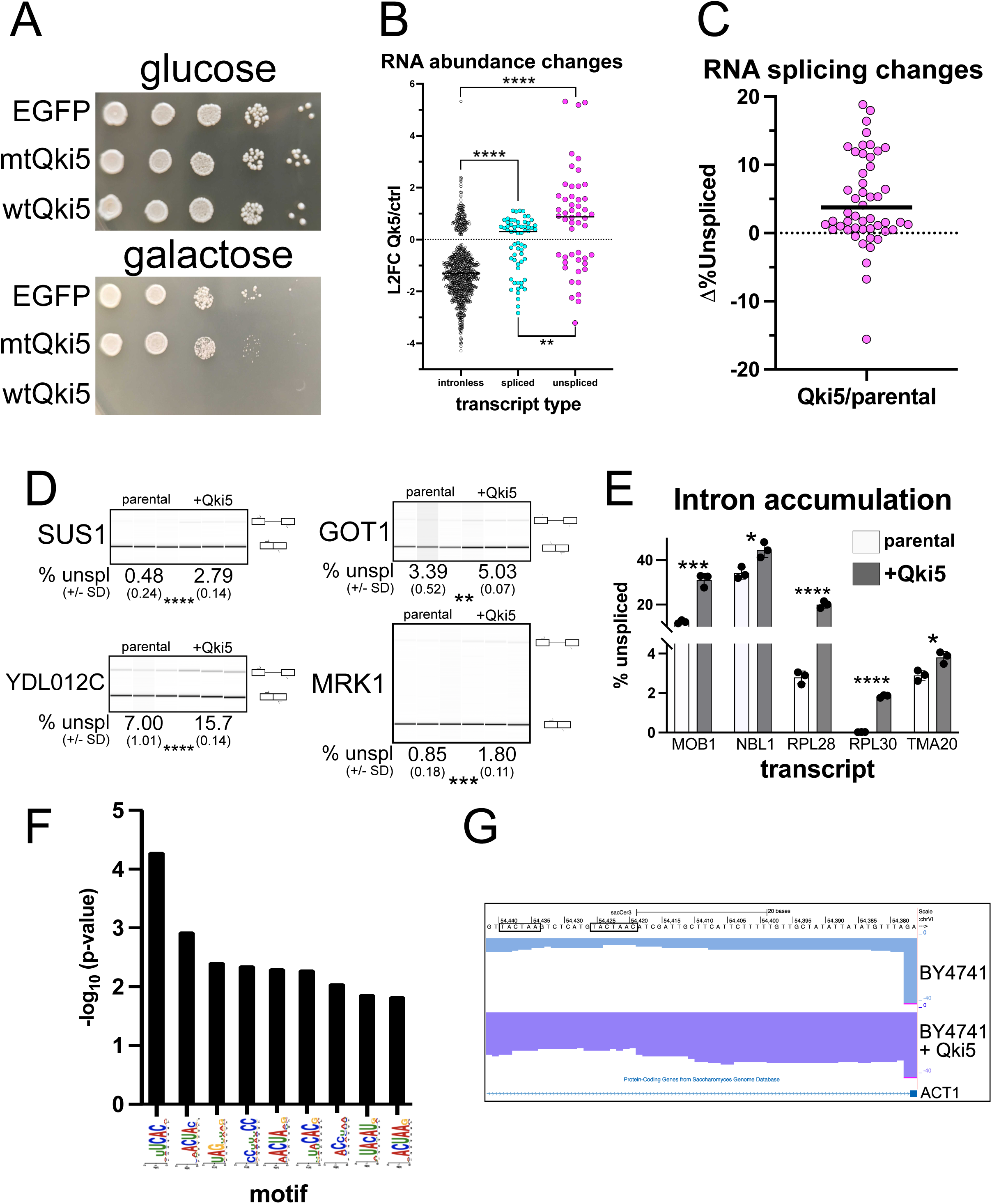
*QKI5* expression is lethal in yeast and represses splicing. A. BY4741 *S. cerevisiae* strain with *EGFP* (top), mutant *QKI5* (mtQKI5; middle), or wild type *QKI5* (wtQKI5; bottom) GAL-inducible transgene cultures grown on either glucose-(top) or galactose-containing (bottom) YPD plates at decreasing densities (left to right) and incubated at 30°C for 72h. B. Changes in RNA abundance measured by Deseq2 analysis of RNA-seq data for intronless, spliced, or unspliced transcripts were measured from RNA extracted from BY4741 cells with *QKI5* expression (induced by galactose-containing media for 4h), or in the BY4147 parental strain also cultured in galactose-containing media for 4h (control), and are shown as L2FC in *QKI5*-induced cells relative to control (n = 3 biological replicates; cutoff *P* < 0.1; abundance cutoff of TPM > 0.2); \*\**P* < 0.01 or \*\*\*\**P* < 0.0001). C. Splicing changes measuring the change in percent unspliced (Δ%Unspliced; y-axis) for BY4741 as described in B.; cutoff *P* < 0.1 by Student’s t-test and base mean > 100. D. RT-PCR with primers that span exon-intron-exon junctions and BioAnalyzer gel-like image showing mean percent unspliced (+/- SD; n = 3) below for introns whose inclusion increased upon *QKI5* ectopic expression and as measured by RNA-seq analysis in C. for the parental or *QKI5*- expressing cells (\*\**P* < 0.01, \*\*\**P* < 0.001, \*\*\*\**P* < 0.0001 by Student’s t-test). E. RT-qPCR analysis measuring mean percent unspliced transcript from RNA extracted from biological triplicate cultures of either the parental control or BY4741 expressing *QKI5* for each transcript shown (+/- SD; \**P* < 0.05, \*\*\**P* < 0.001, \*\*\*\**P* < 0.001). F. Bar graph showing -log_10_ *P*-values (y-axis) of significantly enriched (SEA; *P* < 0.01) motifs observed in 3’ proximal ends of introns whose inclusion increases upon ectopic *QKI5* expression in yeast (x-axis). G. UCSC Genome Browser screen shot showing RNA-seq reads mapping to the *ACT1* transcript intron/exon junction near the 3’ss from RNA extracted from parental control and *QKI5*-induced cells; boxed sequences show two TACTAA elements.

Next we tested how ectopic QKI5 expression impacts the yeast transcriptome. The parental BY4741 strain, or BY4741 with WT QKI5 were grown in galactose-containing media for 4h, and then we collected RNA and performed RNA sequencing and analysis. We used a *S. cerevisiae* genome annotation that specifies intronic regions and pre-mRNAs, spliced mRNAs, or intronless transcripts (76) to map RNA-seq reads to the transcriptome and then measured these different transcript types. We observe higher levels of unspliced pre-mRNA and spliced mRNA when comparing the changes in QKI5 induction to the control, while intronless transcript abundance decreases (cutoff *P* < 0.1; Fig 5B and Supplemental Table S6). Unspliced pre-mRNAs accumulate to a greater degree than spliced mRNAs, supporting the notion that ectopic QKI5 expression perturbs splicing in yeast (\*\**P* < 0.01 by Mann-Whitney U; Fig 5B). We measured changes in splicing by calculating the percentage of unspliced pre-mRNA for each intron-containing transcript (n = 304 (out of 325 total) that passed expression level cutoff of base mean > 100), and discovered that 50 change significantly (cutoff *P* < 0.1 by Student’s t-test; Supplemental Table S7) upon QKI5 expression. Of these, 41 increase in percent unspliced (∼12% of all expressed intron-containing transcripts) while 9 decrease compared to control, which indicates a strong bias toward intron accumulation upon ectopic QKI5 expression (Fig 5C). We validated several of these using RT-PCR (Fig 5D (n = 4) and Supplemental Fig S5C (n = 15)) or RT-qPCR (Fig 5E (n = 5)) and found that each of these instances of repressed splicing upon QKI5 expression that is measured by RNA-seq is also observed using these complementary methods. Notably, the HAC1 intron, which is not removed by the spliceosome (77), shows no increase in intron accumulation upon ectopic QKI5 expression by either RNA-seq or RT-PCR (Supplemental Fig S5C). Next we asked if any intron sequence motifs are enriched that correlate with increased intron retention upon QKI5 expression by obtaining 80 nt of intron sequence upstream of the 3′ss and compared these to 80 nt of intron sequence in 108 control introns that are expressed but unchanged compared to control using SEA ((60) see Methods). Nine motifs are significantly enriched (*P* < 0.01), 4 of which have varying degrees of resemblance to the QKI binding motif (ACUAA) or “half-site” (UAAY) (31, 36, 37). We note that several of these could potentially serve as tandem embedded motifs (for example, U*ACUA<u>A</u>*<u>CUAA</u>C where the first is italicized and the second underlined). A search for these finds that 18% of the introns that accumulated upon QKI5 induction compared to control have either two independent or tandem embedded motifs. In contrast, examination of the introns that are unchanged upon QKI5 induction reveals that only 6% had two intron ACUAA motifs within 80 nt of the 3′ss. The occurrence of two ACUAA elements per intron 3′ss-proximal region is significantly overrepresented in the set of introns that accumulate upon QKI5 ectopic expression (*P* = 0.035 by Chi-squared test), suggesting that these are more sensitive to QKI5 splicing repression. Intriguingly, the essential ACT1 pre-mRNA has dual ACUAA elements in its intron proximal to the 3′ss, like those observed in RAI14 intron 10 (Fig 2A), and its splicing is blocked by QKI5 induction (Fig 5F). Thus, QKI5 expression in yeast is lethal and is due, at least in part, to splicing inhibition.

## Discussion

Our study reveals that QKI can repress the inclusion of alternatively spliced exons by directly competing with SF1/BBP for ACUAA bs. We find that 43% of the alternatively spliced exons that are co-regulated by both SF1 and QKI fall within the “QKI repressed and SF1 activated” category and that these are associated with an ACUAA sequence motif/bs (Figs 1 and 2). We show unambiguously that either of the two ACUAAC elements in RAI14 intron 10 can be used as a bs but that one or the other is required for any splicing, and so must be a bs (Fig 3). Interestingly, deleting either ACUAAC element reduces the level of exon skipping (Fig 3) and QKI5 binding (Fig 4), but increases the level of inclusion (Fig 3) and SF1 binding (Fig 4). Therefore, QKI binds with high affinity to a bona fide dual bs in RAI14 intron 10 to prevent SF1 binding and splicing activation. In budding yeast, where the bs in nearly invariant ACUAAC, ectopic expression of QKI5 blocks pre-mRNA splicing and is lethal (Fig 5). Together these findings demonstrate the validity of the bp competition model by showing these two RBPs directly compete for ACUAA bs substates.

### Consequences of bp competition for pre-mRNA substates and other RNAs

One of the most well-studied examples of competition between RBPs for pre-mRNA substrates is SR proteins that promote exon activation versus hnRNPs that repress exon inclusion (78, 79). This is the predominant manner through which most cell/tissue types carry out alternative pre-mRNA splicing. The model for bp competition between SF1 and QKI that we define here describes a novel but relatively simple pathway through which a specific subset of alternatively spliced exons with ACUAA bs can be regulated. Our findings are consistent with the established role of SF1 in bp recognition and splicing activation, and suggest that QKI is more often a splicing repressor. A nuanced implication of this model is that loss-of-function of one of these proteins leads to a gain-of-function in the other, by relieving the competitive inhibition for a specific number of ACUAA bs that control the inclusion of certain alternatively spliced exons.

How does competition between SF1 and QKI influence transcript fate? In the case of *RAI14* intron 10, the dual ACUAA bs constitute a high-affinity bipartite binding site for dimeric QKI5 protein. In C2C12 myoblasts where QKI5 levels are high, *Rai14* exon 11 is mostly skipped (Fig 2), and this correlates with QKI5 binding (Figs 2 and 4). This binding event appears to promote paraspeckle association, as the paraspeckle proteins NONO, SFPQ, and PSPC1 (74) co-associate with QKI5-bound *RAI14* intron 10 (Fig 4F). This finding is consistent with a recent report that identifies QKI as a paraspeckle-associated protein (75). It is unclear how paraspeckle localization might influence splicing outcomes, but pre-mRNA localization within the nuclear speckle is associated with more efficient splicing (80). It is possible that QKI-directed paraspeckle localization is repressive to pre-mRNA splicing or promotes the ligation of the exons flanking an alternatively spliced one, leading to exon skipping, as we observe in *RAI14*. Further study is required to elucidate how competition between SF1 and QKI for pre-mRNA substates influences their subnuclear localization, splicing, and stability. A recent study suggests that this may be complicated by the fact that QKI itself attenuates paraspeckle biogenesis by promoting the expression of the short *NEAT1* lncRNA isoform, which has lower paraspeckle-promoting activity than the longer isoform (81). It is unclear if SF1 might have a similar role, which is possible, given that QKI promotes expression of the short isoform of NEAT1 via ACUAA elements (to which SF1 might also bind). Therefore, this may constitute a complex feedback loop in which QKI (and/or SF1) can regulate splicing in a paraspeckle-dependent manner but also regulates the degree to which paraspeckles accumulate.

### Branch site competition defines a subset of cell type-specific alternative splicing

Competition for alternatively spliced exons between SR proteins and hnRNPs largely explains the splicing patterns observed in most cell types, but not in the most divergent patterns are observed in muscle or brain cells (82, 83). This is due in part to muscle- or brain-specific RBP expression patterns, and our study uncovers the novel finding that bp competition between SF1 and QKI explains in part how these are enforced. SF1 levels are relatively high in most tissues but can vary by up to 8-fold (59), while QKI levels are much more dynamic. QKI levels are elevated in muscle, but undetectable in most neurons or predominated by the cytoplasmic isoforms QKI6 and QKI7 in glia (45, 46, 48, 49, 84). Therefore, in muscle, the nuclear QKI:SF1 ratio is high, whereas in many brain cell types, it is low (or zero), resulting in opposite patterns of isoform-specific transcriptomes and proteomes within the context of the bs competition model. This is also relevant in developmentally regulated systems, where QKI5 levels increase during cardiac cell differentiation (85) but decrease as neural stem cells commit to mature cell types (39). Similarly, when embryonic stem cells exit pluripotency and commit to the endoderm lineage, QKI levels decrease; however, QKI levels increase upon commitment to the mesodermal lineage, and a QKI-associated splicing gain-of-function is observed (86). On the other hand, SF1 reduction, leading to increases in unspliced RNA and skipped exons, has been observed in a model of aging and correlates with age-related decline in fitness (87). In summary, bs competition between SF1 and QKI appears to be a novel molecular mechanism through which different cell type/tissue-, lineage-, developmental-specific, and homeostatic splicing programs can be achieved and regulated.

### Evolutionary implications of bp competition

Yeast and other single-celled eukaryotes lack *QKI*, but multicellular eukaryotes and plants have the gene or an orthologous one. Strikingly, we found that 12% of the introns expressed in yeast strain BY4741’s splicing was inhibited by ectopic QKI5 expression, and that these cells developed a large-budded phenotype due to a failure to divide (Supplemental Fig S5A and S5B). This phenotype has been observed due to lack of *TUB1* pre-mRNA splicing (88), although we did not observe mis-splicing of it (or *TUB3*) in our study. The introns most sensitive to QKI5 have evidence of the “bipartite” QKI motif, which is consistent with high-affinity binding of dimeric QKI (31) and similar to that observed in RAI14 intron 10. It is interesting that yeast lack *QKI* and do not exhibit “alternative splicing” per se, and also have a more invariant bs. This is UACUAAC in *S. cerevisiae* (14, 16, 19), but *Schizosaccharomyces pombe* have a slightly more degenerate bs (89), and its introns share some features with both budding yeast and mammals (90, 91). Nevertheless, the consensus bs in the latter still represents what would be a strong QKI binding motif (31, 36), suggesting that QKI expression might also be lethal. Thus, it would be evolutionarily unfavorable to select for *QKI* or an ortholog/close relative.

The metazoan STAR family of RBPs, with the exception of SF1/BBP which is the most divergent member, all contain the QUA1 dimerization domain (30). In the case of QKI (or ASD2 in *C. elegans* (92) or HOW in *Drosophila* (93)), the binding motif ACUAA, along with the presence of a YAAY “half site” within about 20 nt, requires more specificity and generates additional points of contact on an RNA, which may explain in part its higher binding affinity than SF1. Perhaps it would not have been evolutionarily advantageous for the majority of bs in multicellular organisms to be UACUAAC/high-affinity QKI motifs (fewer than 20% of human introns have this bs (94)). So more degenerate bs were selected for in metazoans. However, the presence of some ACUAA bs could help diversify the function of cell type-specific protein isoforms. For example, many cytoskeletal/contraction-related protein isoforms are specific to muscle (95, 96), and many of these are alternatively spliced exons are repressed by QKI through binding UACUAAY in introns upstream of them (38, 40, 41, 85). Similar observations in worms and flies suggest the intriguing possibility that the *QKI* gene and the more extensive bs degeneracy that is observed in metazoans and plants may have co-evolved. One way this would have been executed, but also constrained, is through bs competition between QKI and SF1/BBP. Consequently, this may have allowed for expansion of the transcriptome and proteome through the diversification of alternative splicing.

## Materials and Methods

### Cell culture

C2C12 mouse myoblasts and HEK293 cells were cultured in Dulbecco’s Modified Eagle Medium (DMEM) supplemented with high glucose (Invitrogen) and 10% (v/v) heat-inactivated fetal bovine serum (Thermo Fisher). Cells were maintained at 37°C in a humidified atmosphere with 5% CO_2_. The HEK293 QKI KO cells were generated as previously described (97, 98).

### Plasmids and transfections

The pDUP51 splicing reporter plasmid (99) was used as a backbone to generate the RAI14 exon 11 plasmid. A 243 bp fragment upstream of exon 11, exon 11, and a 95 bp fragment downstream of the exon were PCR-amplified from H9 human embryonic stem cell genomic DNA; the forward primer contained an ApaI site and the reverse primer contained a BglII site. These and the pDUP51 plasmid were digested with ApaI and BglII (NEB), then ligated into pDUP51 at the ApaI and BglII restriction sites. The resulting pDUP-RAI14 exon 11 plasmid was confirmed by Sanger sequencing. Mutant constructs were generated using the Q5® Site-Directed Mutagenesis Kit and were also verified by Sanger sequencing. The myc:QKI5 plasmids used were described previously and were a kind gift from Sean Ryder (38, 86). Generation of the pcDNA3.1-tdTomato plasmid was also previously described (38). The pcDNA3.1-SF1 plasmid was generated by Gibson Assembly using cDNA from H9 human embryonic stem cells as a template for PCR amplification and confirmed by Sanger sequencing.

Transfections were carried with Lipofectamine 2000 (Invitrogen) using a “in tube transfection” protocol, where 2.5X10^5^ cells were added to a tube containing Lipofectamine and the appropriate volume of reagent, following the manufacturer’s instructions, with 100 ng DNA mix in Gibco Opti-MEM (Thermo) or 30 pmol of siRNA, incubated for 20 minutes at RT, plated in 12 well plates containing DMEM 10% FBS and incubated overnight. Cells were harvested 24 hours after transfection.

For the RNA affinity chromatography, the j6f1 aptamer under the control of T7 RNA promoter was obtained by the digestion of the vector pcDNA5-aptamer plasmid, and was a gift from Chloe Nagasawa (University of Texas Medical Branch, Galveston, Texas, US) from the lab of Mariano Garcia-Blanco (University of Virginia). RAI14 sequence consisting of 60 bp fragment of RAI14 intron 10 upstream RAI14 exon 11 was obtained from a gBlock (IDT), digested with HindIII and Not1 restrictions enzymes and cloned into the digested PCNA5 aptamer plasmid.

For yeast transformation the plasmid pJW1666 (gift from Jonathan Weissman (Addgene plasmid # 112040; http://n2t.net/addgene:112040; RRID:Addgene_112040(100)) was used to clone the WT or mutant QKI sequences, or the GFP present on WT the plasmid was used as control.

### RNA extraction, RT-PCR and RT-qPCR

RNA was extracted from cells using Trizol (Thermo Fisher). For direct extraction from cell plates, Trizol was added to the wells and samples were vortexed. For extraction from cell lysates, cells were extracted with RSB100 (100mM Tris-HCl pH 7.4, 0.5% (v/v) NP-40, 0.5% (v/v) Triton X-100, 0.1% (w/v) SDS, 100 mM NaCl and 2.5 mM MgCl2). Samples were vortexed, chloroform was added then the samples were incubated at RT for 2 minutes. After centrifugation at 13,000 RPM for 15 minutes at 4°C, the aqueous phase was transferred to a new tube and samples were processed with the ReliaPrep RNA MiniPrep system (Promega), according to the manufacturer’s instructions. Reverse transcription was performed with iScript™ Reverse Transcription Supermix (BioRad) according to manufacturer instructions. RT-PCR was performed using Taq2x Master Mix (NEB), and cycle numbers were determined empirically to prevent over-amplification. PCR products were analyzed by agarose gel and via Agilent 2100 BioAnalyzer.

For RT-qPCR, cDNA was diluted 1:8 and used with Applied Biosystems™ SYBR™ Green Universal Master Mix on a Step One Plus Real-Time PCR machine (Applied Biosystems). RT-qPCR cycling conditions were: 95°C for 10 minutes, [95°C 15 seconds, 60°C 60 seconds] (40 cycles). DUP-RAI14 exon 11 reporter RNA abundances were analyzed by calculating the ΔCt values normalized to EEF1A1. The ΔΔCt values were then calculated as 2^-ΔCt for each sample. The average ΔΔCt of the wild-type reporter was used as a normalization factor to determine the relative RNA abundance in the mutant reporters.

### Western blotting

Proteins were extracted using RSB100 (100 mM Tris-HCl pH 7.4, 0.5% (v/v) NP-40, 0.5% (v/v) Triton X-100, 0.1% (w/v) SDS, 100 mM NaCl and 2.5 mM MgCl_2_) with EDTA-free protease inhibitor cocktail (SIGMAFAST, Sigma) and 1 mM PMSF. The buffer was added directly to the wells of the plates, scraped, collected, and kept on ice, while vortexing every five minutes for 30 minutes, followed by centrifugation at 14000 RPM for 15 minutes at 4°C. The supernatant was transferred to a new tube. Samples were stored at −80°C until use.

Protein concentration was measured using the Bradford assay (BioRad (101)). The same amount of protein from each sample was loaded onto 10% SDS-PAGE (typically 15-30 µg) and then transferred to a 0.45 µm nitrocellulose membrane (Thermo). The membrane was blocked (with 5% non-fat dry milk dissolved in tris-buffered saline (TBS)) for one hour at room temperature with constant agitation. Primary antibody incubations were performed overnight at 4°C with constant agitation in TBS 0.01% (v/v) tween (TBS-T) with 5% non-fat dry milk using the following antibodies: Anti-Pan-QKI Antibody, clone N147/6 (MABN624, Sigma-Aldrich) (1:2000), IgM-anti-GAPDH (G8795, clone GAPDH-71.1, Sigma-Aldrich) (1:40000), α/β-Tubulin (#2148, Cell Signaling) (1:5000), anti-SF1 (#A303-213a, Bethyl Laboratories) (1:5000), and anti-HTATSF1 (# 20805-1-AP, Thermofisher) (1:2000), all diluted in TBS containing 5% (w/v) milk and 0.01% (v/v) Tween-20 (TBST). The following day, the membranes were washed three times with TBST with 5% (w/v) milk at RT. Secondary infrared conjugated antibodies: IRDye 800CW Goat anti-Mouse IgG2b (P/N 926-32352), IRDye 680RD Goat anti-Mouse IgM (P/N: 926-68180), IRDye 800CW Goat anti-Rabbit IgG (P/N: 926-32211) and IRDye 680RD Goat anti-Rabbit IgG (P/N: 926-68071) from Li-Cor were diluted according to manufacturer instructions (1:15,000 for 800CW or 1:20,000 for 680RD/LT conjugates) in TBST with 5% (w/v) non-fat dry milk for 1 hour with constant agitation. Membranes were washed 3 times every 5 minutes before visualization on Odyssey CLx imager (Li-Cor).

### Yeast transformation

Yeast transformation was performed as described by Ito *et al* with modifications (102) using the BY4741 strain. A 50 mL of culture of BY4741 was grown to a density of 5-10^6 overnight. Following centrifugation, the pellet was resuspended in 1ml of deionized water, centrifuged at 12000 RPM and resuspended in 0.5ml of 1X TE-LiAc (100mM tris-HCl, 10mM EDTA pH 7.5; LiAc pH 7.5). Subsequently, cells were resuspended in 3x volume of 1X TE-LiAc and incubated at 30°C with agitation. 2 µg of purified PCR of WT QKI, mutant QKI and GFP plasmids and 7 µL of salmon sperm carrier DNA were added to 200 µL of the yeast cells and the mixture was incubated for 45 min at 30°C with agitation. The following steps included a 3 h incubation at 30°C with agitation, a 20-minute heat shock at 42°C, centrifugation at 12000 RPM, and resuspension in YEPD followed by a 45-minute incubation. After a 12000 RPM centrifugation, cells were resuspended in water and spread in URA-selection plates, then incubated at 30° C until transformants appeared.

### Yeast genomic DNA extraction

Genomic DNA extraction was performed using the method developed by Klassen and collaborators with a few modifications (103). Cells were patched on URA-selection plates, and 50 µL of cell volume was scraped off the plate and resuspended in zymolase 20T 8 mg/ml. After vortexing the samples, they were incubated at 37°C for 1 hour. After incubation, the cells were spun down, the supernatants were aspirated, and the cells were resuspended in 250 µL of 10% SDS. Then, they were vortexed and incubated for 30 minutes at 65°C. After incubating the samples on ice for 5 minutes, 100 µL of 5 M potassium acetate, pH 7.5, was added, and the samples were incubated on ice for 1 hour. Tubes were spun at 14000 RPM for 15 minutes, and the supernatant was transferred to a new tube. Then 400 µL of ice-cold 95% EtOH was added, vortexed briefly, and spun for 15 minutes at 14,000 RPM. The supernatants were decanted, and the DNA pellet was dried at 30°C for 15 minutes. It was then dissolved in 100 µL of ultra-pure water and placed in a mixer for 20 minutes at 2000 RPM. The extracted genomic DNA was used to perform PCR with high-fidelity polymerase (Takara Bio Inc) to confirm the transgene expression by Sanger sequencing.

### Yeast RNA extraction

RNA was extracted from yeast cells as previously described (104). Briefly, 1 mL of yeast culture was pelleted and then resuspended in 400 µLof AE buffer (50 mM, pH 5.2 of NaOAc, 10Mm EDTA). After the addition of 40 µL of 10% SDS and 400 µL of PCA, the samples were incubated at 65°C for 10 minutes. After a 5-minute incubation on ice, samples were placed in phase lock gel tubes, centrifuged at 14,000 RPM for 5 minutes, and centrifuged again using the same conditions following another chloroform addition; this chloroform wash was repeated, and the samples were centrifuged one final time. The aqueous phase was transferred to a new 1.5 mL tube, 50 µL of 3M sodium acetate, pH 5.2, was added, followed by 2 volumes of 100% ethanol. Samples were centrifuged at 14,000 RPM for 15 minutes, the ethanol was removed, and the pellet was washed with 70% ethanol. Then, the samples were centrifuged at 14,000 RPM for 5 minutes, the ethanol was removed, and the samples were air-dried. The pellet was resuspended in RNAse-free water and then vortexed. To remove any potential DNA contamination, 1 µg of RNA was treated with DNase Turbo following the manufacturer’s instructions. RNA was extracted after DNase treatment with phenol/chloroform/isoamyl alcohol and then ethanol precipitation. RNA concentration was measured using the NanoDrop (ThermoFisher).

### Yeast spot assay

One milliliter of freshly harvested cells grown overnight in 10mL of YEPD were centrifuged, washed in ultra-pure water and sonicated. The cells were then diluted to a concentration of 1×10^7^. Serial ten fold (up to 5 times) were prepared on both Galactose (Gal) and non-galactose plates. The plates were incubated at 30° C for 3 days.

### Yeast growth curve

Yeast was grown in YEPD overnight. When the OD660 reached approximately 0.5, transgene activation was induced by adding galactose dissolved in water to a final concentration of 2% or water as control. For each time point, samples were collected for cell count and RNA extraction.

### RNA sequencing and analysis

RNA-seq data from the ENCODE project for shQKI and shNT or shSF1 and shNT (57, 105) were downloaded from GEO and analyzed as previously described (86) using Vast-tools (58, 59) or rMATS (61) to compare alternative splicing observed in shQKI relative to shNT, or shSF1 relative to shNT

The subsequent Vast-tools data files were filtered by enlisting a cutoff of dPSI > |10| and MVdPSI95 > 0 to measure alternatively spliced events that changed significantly upon either QKI or SF1 knockdown compared to the control. For “co-regulated” exons, the above cutoff was required for either shQKI compared to control or shSF1 compared to control, but in both datasets the MVdPSI95 had to be > 0. rMATS analysis and rMAPS2 (62, 63) motif enrichments were done using default conditions and with custom motifs added to measure various bp-like and potential SF1 binding motifs (ACT[ACTG]AG, [ACTG]CT[AG][CT], TAA[CT], TAA[CT]T[ACTG]A[CT], TACTAAC, TACTAA, ACTAA[CT], TACTAA[CT], CTAAC[ACG]).

Strand-specific RNAseq libraries were prepared from 1 µg of yeast total RNA using NEBNext poly(A) mRNA Magnetic Isolation module (NEB, E7490) and NEBNext Ultra II Directional RNA Library Prep kit for Illumina (NEB, E7760) following the manufacturer’s recommended procedure. The six libraries were pooled in equal molar concentration and sequenced on Illumina NextSeq 550 for PE 150 base pair sequencing, yielding about 30M paired-end reads each. The associated datafiles have been uploaded to GEO under accession GSE273838. The RNAseq reads were filtered for low-quality bases and trimmed of adapter sequences using trimmomatics (v.0.39). The trimmed reads were aligned to the yeast reference genome SacCer3 and a custom yeast annotation file, which accounts for all yeast introns (46) (and allows measurement of unspliced, spliced, and intronless transcripts) using kallisto (v.0.50) to generate the abundance.tsv file for either normalized (TPM) or total (counts) reads for each sample. The TPM file was used as input for Deseq2 analysis (v1.42.1) to identify changes in transcript abundance (abundance cutoff TPM > 0.2 and significance cutoff *P* > 0.1; Supplemental Table S6), and the read counts file was used to calculate the percentage of unspliced RNA (percent unspliced = (unspliced/(unspliced+spliced))*100) for each transcript that was expressed (base mean > 100) and statistical significance was measured using the Student’s t-test (double check w/Jose) with a change in percent unspliced *P* < 0.1 considered significant (Supplemental Table S7).

Motif analysis of both the yeast and HepG2 RNA-seq dataset were performed using SEA (60) and the former used 80 nt of intron sequence upstream of the 3′ss of yeast introns, 38 of which increased (*P* < 0.1) upon ectopic QKI5 expression compared to the parental control (this represents 38 of the 41 we observed that increased in inclusion upon QKI5 expression, as the other two were located on ChrM and did not overlap with “Talkish Standard Introns” (76) which was a requirement for out motif analysis), compared to a background set of 106 introns that were detectable (base mean > 100) but unchanged (P > 0.2, change in percent unspliced < |1|) upon ectopic QKI5 expression compared to the parental control. For the HepG2 intron set, we extracted the intron sequence that began 20 nt upstream of the 3′ss (to exclude analyzing potential differences in 3′ss or pY tracts) and spanned 60 nt upstream of the 3′ss of introns that were co-regulated by QKI and SF1 (dPSI > |10| and MVdPSI at 95% confidence interval > 0 in either dataset or the other and at least MVdPSI at 95% confidence interval > 0 in both datasets, as determined by Vast-tools) and performed SEA, and then also performed an identical analysis but using 1000 introns that were detectable (base mean > 100) but unchanged in either shQKI relative to shNT or shSF1 relative to shNT (dPSI < |1|, MVdPSI = 0). In both cases, the motif set that was interrogated was from Ray et al 2013 (106).

### RNA affinity chromatography

Tobramycin RNA affinity chromatography was performed as previously described (65) with several minor modifications. Briefly, the aptamer and RAI14 WT and mutant RNAs were produced after the vectors were digested with NotI (NEB) using the HiScribe T7 High Yield RNA Kit (NEB). Then 120 picomoles of RNA of the aptamer and RAI14 mutants, and 200 picomoles of wt RAI14 were heated in RNA binding buffer (20 mM Tris-HCl, 1 mM CaCl2, 1 mM MgCl2, 300 mM KCl, 0.1 mg/ml tRNA, 0.5 mg/ml BSA, 0.01% (v/v) NP-40, 0.2 mM DTT) at 95 °C for 5 minutes and transferred to RT for 30 minutes. Samples were incubated with 60 µL of tobramycin-coupled sepharose matrix at 4°C for 2 hours with head-over-tail rotation. The beads where washed three times with washing buffer (20 mM Tris-HCl, 1 mM CaCl2, 1 mM MgCl2, 145 mM KCl, 0.1% (v/v) NP-40, 0.2 mM DTT) and incubated with a 285 µL solution of 32% of C2C12 NE (as previously described (107)), 32 mM KCL, 2 mM MgCl2, 2 mM ATP and 20 mM creatine phosphate (CP) (the reactions were also performed without MgCl2, ATP and CP in the NE solution) for 7.5, 15 and 30 min at 30°C with head over tail rotation. After NE incubation, the matrix was washed three times with a higher salt concentration of washing buffer (150 mM KCl). The beads were eluted with 5 mM of tobramycin in 20 mM Tris-HCl, 1 mM CaCl2, 1 mM MgCl2, 145 mM KCl, 2 mM MgCl2, 0.2 mM DTT in a 125 µL solution. The protein eluate was precipitated with acetone and suspended in LDS sample buffer for WB and in 5% (w/v) SDS, 50 mM TEAB pH 7.1 for MS analysis.

### Protein precipitation

Acetone was used to precipitate the proteins from the RNA affinity chromatography-eluted samples. Four times the sample volume of cold acetone was added to each sample and vortexed. Samples were incubated for 60 minutes at −20°C for one hour, following a 10 minutes centrifugation at 14,000 RPM at 4C. The acetone was decanted, and the samples were air-dried for 15 minutes. Pellets were resuspended with 1x Laemmli buffer for western blot or with 5% (w/v) SDS, 50 mM TEAB, pH 7.1, for liquid chromatography with tandem mass spectrometry.

### Protein digestion

The samples were prepared as previously described (108). Briefly, 25 µg of protein from the above were reduced with 10 mM Tris(2-carboxyethyl) phosphine (TCEP) (77720, Thermo) and incubated at 65 °C for 10 min. The sample was then cooled to room temperature and 1 μL of 500 mM iodoacetamide acid was added and allowed to react for 30 min in the dark. Then, 3.3 μL of 12% phosphoric acid was added to the protein solution followed by 200 uL of binding buffer (90% Methanol, 100 mM TEAB pH 8.5). The resulting solution was added to S-Trap spin column (protifi.com) and passed through the column using a bench top centrifuge (60s spin at 1,000 x *g*). The spin column is washed with 150 µL of binding buffer and centrifuged. This is repeated twice. 30 µL of 20 ng/ µL trypsin is added to the protein mixture in 50 mM TEAB pH 8.5, and incubated at 37^○^C overnight. Peptides were eluted twice with 75 µL of 50% acetonitrile, 0.1% (v/v) formic acid. Aliquots of 20 µL of eluted peptides were quantified using the Quantitative Fluorometric Peptide Assay (Pierce, Thermo Fisher Scientific). Eluted volume of peptides corresponding to 5.5 µg of peptides are dried in a speed vac and resuspended in 27.5 uL 1.67% (v/v) acetonitrile, 0.08% (v/v) formic acid, 0.83% (v/v) acetic acid, 97.42% water and placed in an autosampler vial.

### Data dependent acquisition NanoLC MS/MS Analysis

Peptide mixtures were analyzed by nanoflow liquid chromatography-tandem mass spectrometry (nanoLC-MS/MS) using a nano-LC chromatography system (UltiMate 3000 RSLCnano, Dionex), coupled on-line to a Thermo Orbitrap Fusion mass spectrometer (Thermo Fisher Scientific, San Jose, CA) through a nanospray ion source (Thermo Scientific) similar to as we have described previously (109). A trap and elute method was used. The trap column was a C18 PepMap100 (300 µm X 5mm, 5 µm particle size) from ThermoScientific. The analytical column was an Acclaim PepMap 100 (75 µm X 25 cm) from (Thermo Scientific). After equilibrating the column in 98% solvent A (0.1% formic acid in water) and 2% solvent B (0.1% (v/v) formic acid in acetonitrile (ACN)), the samples (2 µL in solvent A) were injected onto the trap column and subsequently eluted (400 nL/min) by gradient elution onto the C18 column as follows: isocratic at 2% B, 0-5 min; 2% to 24% B, 5-86 min; 24% to 44% B, 86-93 min; 44% to 90% B, 93-95 min; 90% B for 1 minute, 90% to 10% B, 96-98 min; 10% B for 1 minute 10% to 90% B, 99-102 min 90% to 4% B; 90% B for 3 minutes; 90% to 2%, 105-107 min; and isocratic at 2% B, till 120 min.

All LC-MS/MS data were acquired using XCalibur, version 2.5 (Thermo Fisher Scientific) in positive ion mode using a top speed data-dependent acquisition (DDA) method with a 3 sec cycle time. The survey scans (*m/z* 350-1500) were acquired in the Orbitrap at 120,000 resolution (at *m/z* = 400) in profile mode, with a maximum injection time of 100 msec and an AGC target of 400,000 ions. The S-lens RF level is set to 60. Isolation is performed in the quadrupole with a 1.6 Da isolation window, and CID MS/MS acquisition is performed in profile mode using rapid scan rate with detection in the ion-trap, with the following settings: parent threshold = 5,000; collision energy = 32%; maximum injection time 56 msec; AGC target 500,000 ions. Monoisotopic precursor selection (MIPS) and charge state filtering were on, with charge states 2-6 included. Dynamic exclusion is used to remove selected precursor ions, with a +/- 10 ppm mass tolerance, for 15 sec after acquisition of one MS/MS spectrum.

### DDA Database Searching

Tandem mass spectra were extracted and charge state deconvoluted by Proteome Discoverer (Thermo Fisher, version 2.2.0388). Charge state deconvolution and deisotoping were not performed. All MS/MS samples were analyzed using Sequest (Thermo Fisher Scientific, San Jose, CA, USA; in Proteome Discoverer 2.5.0.402). Sequest was set up to search the conanical mouse proteome and a contaminant database cRAP assuming the digestion enzyme trypsin. Sequest was searched with a fragment ion mass tolerance of 0.60 Da and a parent ion tolerance of 10.0 PPM. Carbamidomethyl of cysteine was specified in Sequest as a fixed modification. Deamidated of asparagine and glutamine, oxidation of methionine and acetyl of the n-terminus were specified in Sequest as variable modifications.

### Criteria for Protein Identification

Scaffold (version Scaffold_4.11.1, Proteome Software Inc., Portland, OR) was used to validate MS/MS based peptide and protein identifications. Peptide identifications were accepted if they could be established at greater than 95.0% probability by the Scaffold Local FDR algorithm. Protein identifications were accepted if they could be established at greater than 99.0% probability and contained at least 2 identified peptides. Protein probabilities were assigned by the Protein Prophet algorithm (110). Proteins that contained similar peptides and could not be differentiated based on MS/MS analysis alone were grouped to satisfy the principles of parsimony. Proteins sharing significant peptide evidence were grouped into clusters. The resulting normalized spectral counts from the input NE, each RAC substrate (WT, upDEL, dnDEL, 2xDEL, and APT Only control) were obtained, and those values were used to calculated background-corrected levels of enrichment relative to NE ((NSC_RAC_ – NSC_APT_)/NSC_NE_). We used those calculations to construct the heatmap shown in Fig 4B and generate the counts shown in Fig 4C, by including proteins with experimental LC-MS/MS evidence of being members of early spliceosomes (E-complex or 17S U2 snRNP (66–68)), and that had a positive value of background-corrected levels of enrichment relative to NE in at least one of the 4 RAC substrates tested. See Supplemental Table S4.

### Data independent acquisition NanoLC MS/MS Analysis

Peptide mixtures were analyzed by nanoflow liquid chromatography-tandem mass spectrometry (nanoLC-MS/MS) using a nano-LC chromatography system (UltiMate 3000 RSLCnano, Dionex), coupled on-line to a Thermo Orbitrap Eclipse mass spectrometer (Thermo Fisher Scientific, San Jose, CA) through a nanospray ion source. A direct injection method is used onto an analytical column; Aurora (75um X 25 cm, 1.6 µm) from (ionopticks). After equilibrating the column in 98% solvent A (0.1%(v/v) formic acid in water) and 2% solvent B (0.1% (v/v)formic acid in acetonitrile (ACN)), the samples (2 µL in solvent A) were injected (300 nL/min) by gradient elution onto the C18 column as follows: isocratic at 2% B, 0-10 min; 2% to 27% 10-98 min, 27% to 45% B, 98-102 min; 45% to 90% B, 102-103 min; isocratic at 90% B, 103-104 min; 90% to 15%, 104-106 min; 15% to 90% 106-108 min; isocratic for two minutes; 90%-2%, 110-112 min; and isocratic at 2% B, till 120 min.

All LC-MS/MS data were acquired using an Orbitrap Eclipse in positive ion mode using a data-independent acquisition (DIA) method with a 16Da windows from 400-1000 and a loop time of three seconds. The survey scans (m/z 350-1500) were acquired in the Orbitrap at 60,000 resolution (at m/z = 400) in centroid mode, with a maximum injection time of 118 msec and an AGC target of 100,000 ions. The S-lens RF level was set to 60. Isolation was performed in the quadrupole, and HCD MS/MS acquisition was performed in profile mode using the orbitrap at a resolution of 30000 using the following settings: collision energy = 33%, IT 54ms, AGC target = 50,000. These conditions were duplicated to create six gas-phase fractions of the NE sample using 4Da fully staggered windows in 100 m/z increments from 400-1000 m/z, as described (111).

### DIA Database Searching

The raw data was demultiplexed to mzML with 10 ppm accuracy after peak picking in MSConvert (112). The resulting mzML files were searched in MSFragger (113) and quantified via DIA-NN (https://github.com/vdemichev/DiaNN) using the following settings: peptide length range 7-50, protease set to Trypsin, 2 missed cleavages, 3 variable modifications, clip N-term M on, fixed C carbamidomethylation, variable modifications of methionine oxidation and n-terminal acetylation, MS1 and MS2 accuracy set to 20 ppm, 1% FDR, and DIANN quantification strategy set to Robust LC (high accuracy). The files were searched against a database of human acquired from Uniprot (18^th^ December, 2023). The gas-phase fractions were used only to generate the spectral library, which was used for analysis of the individual samples.

Statistical analysis was performed using Fragpipe-Analyst (114) using an R script based on the ProteinGroup.txt file produced by DIA-NN. First, contaminant proteins, reverse sequences and proteins identified “only by site” were filtered out. In addition, proteins identified by only a single peptide and those not consistently identified or quantified in the same condition are also removed. The DIA data was converted to log_2_ scale, samples were grouped by conditions, and missing values were not imputed. Protein-wise linear models combined with empirical Bayes statistics were used for the differential expression analyses. The limma package from R Bioconductor was used to generate a list of differentially expressed proteins for each pair-wise comparison. A cutoff of the adjusted *P*-value of 0.05 (Benjamini-Hochberg method) along with an absolute log_2_ fold change of 1 has been applied to determine significantly regulated proteins in each pairwise comparison. We also used the E/U2 list to focus our analysis on early spliceosome proteins, and expanded this to include an RBP list (115) to assess non-spliceosome-annotated RBP interactions. The E/U2 or RBPs shown in Fig 4E were included if 1) spectral counts were detected in both the numerator and denominator RAC substrates, 2) log2 fold change ≥ |0.2| and *P*-value was < 0.01 in at least one of the three comparison groups, and 3) data were present for each comparison (no zero or undetectable values). See Supplemental Table S5.

## Acknowledgements

We thank Manuel Ares Jr. (UCSC) for helpful discussion regarding the experiments in *S. cerevisiae* and for piloting an early version of the experiment. We thank Louise Prakash (UTMB) for allowing us to use equipment that made it possible for us to perform the experiments in *S. cerevisiae*, and Robert E. Johnson of the Parakash lab for technical assistance. We thank Steve Widen (UTMB, retired) for technical assistance with the QKI and SF1 knockdown RNA-seq datasets. We thank Mariano Garcia-Blanco and Chloe Nagasawa from Mariano’s lab for discussion and technical suggestions with RNA affinity chromatography. We also thank Eric Van Nostrand (Baylor College of Medicine) and Manuel Ares Jr. for pre-submission review of the manuscript.

## Author’s Contributions

KLPC, WKR, REJ, and WSF designed and optimized experimental approaches; KLPC and REJ performed experiments; KLPC, JA, WSF analyzed data; HH performed upstream bioinformatic analysis of the RNA-seq data and JA, JPD, and WSF performed downstream analyses; WKR and WSF analyzed proteomics data; KCL and WSF provided reagents; WSF supervised the study; WSF funded the study; WSF wrote the paper; all authors have agreed to the final version of the manuscript.

---

**Supplemental Figure S2:**
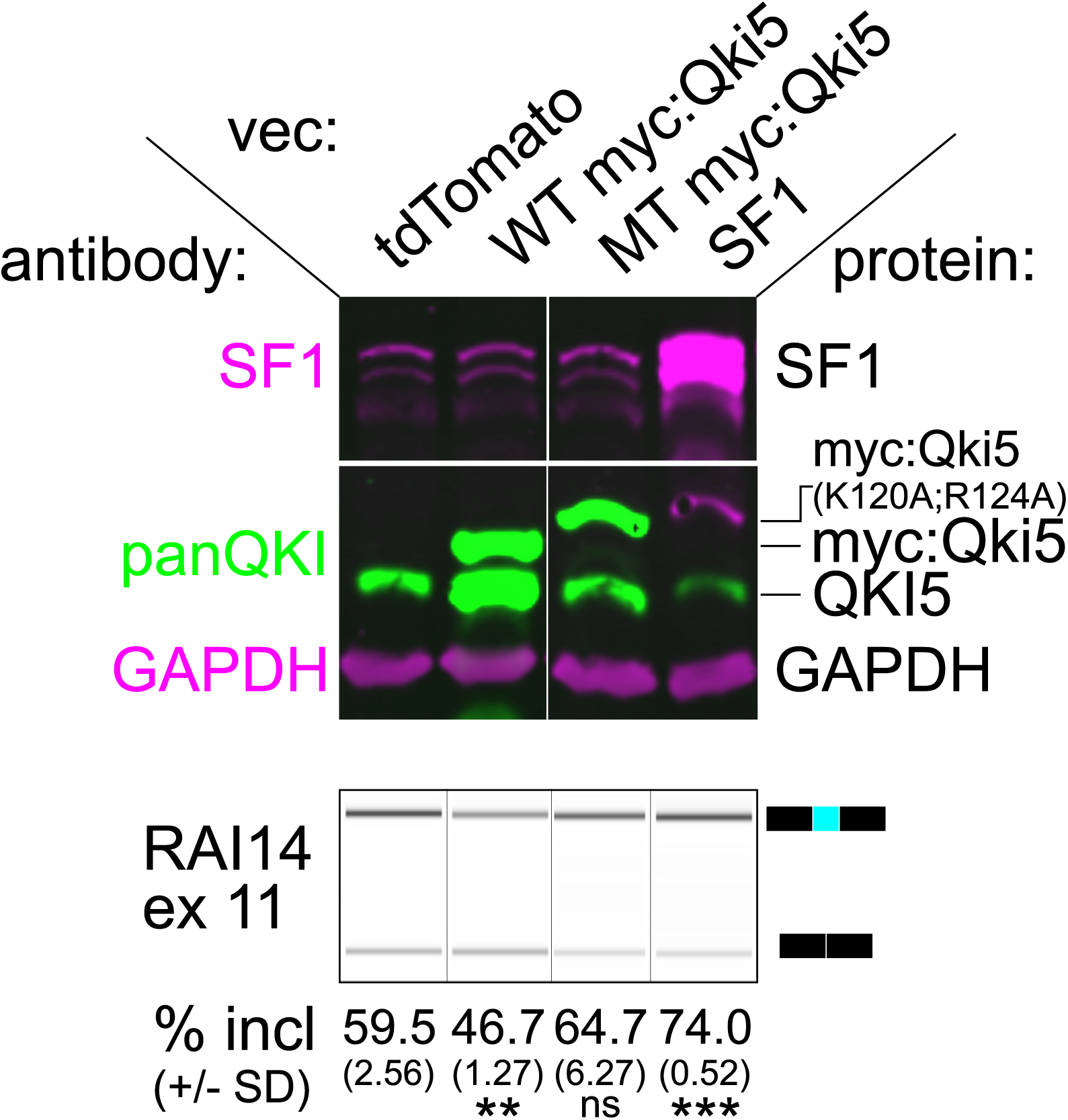
Western blot and RT-PCR of proteins and RNA extracted from WT HEK293 cells transfected with tdTomato, WT myc:Qki5, MT myc:Qki5 and SF1. The top panel shows a western blot probed with anti-SF1 (magenta), anti-PanQKI (green, middle) and anti-Gapdh (magenta, bottom). Below RT-PCR products analyzed on a Bioanalyzer from RNA extracted from transfected WT HEK 293 cells with mean percent included and ± standard deviation bellow (\*\**P* < 0.01,\*\*\**P* < 0.001 by Student’s t-test). The results shown are representative of 3 biological replicates.

**Supplemental Figure S3:**
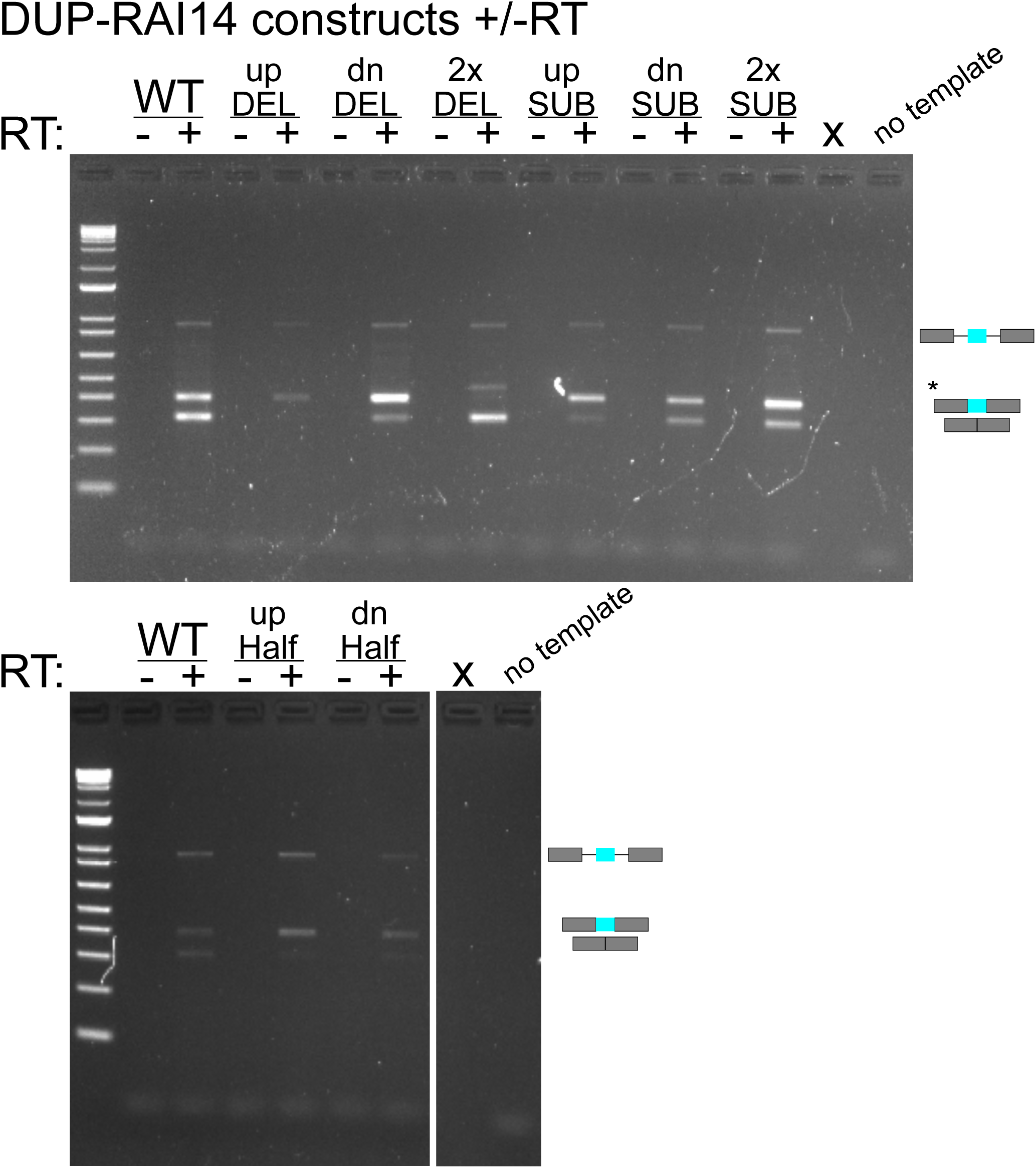
Agarose gel showing PCR amplification in the absence (-) or presence (+) of reverse transcriptase. PCR was performed with RNA from C2C12 cells transfected with *RAI14* reporter plasmids.

**Supplemental Figure S4:**
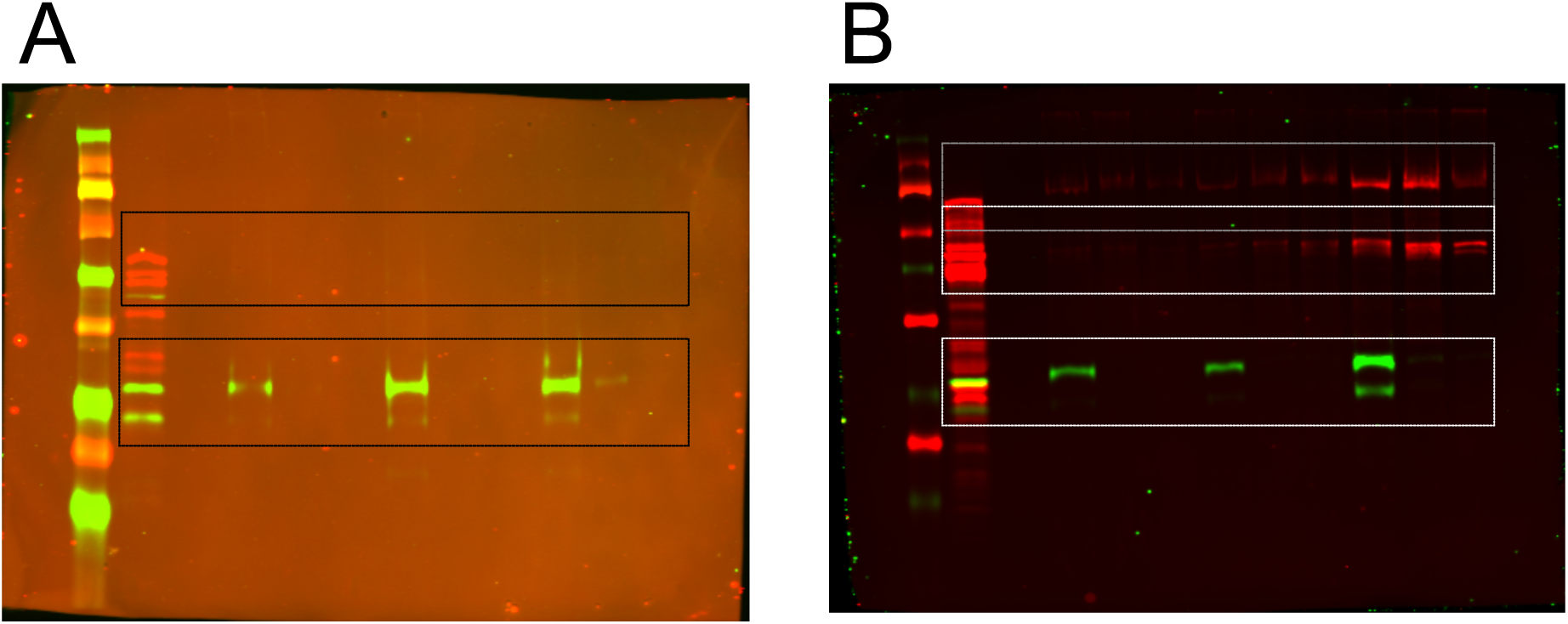
Uncropped western blots from Fig 4. A. Shows uncropped western blot image that is used to construct Fig 4D; QKI is shown in green and SF1 in red and the boxes indicate the cropped region. B.

**Supplemental Figure S5:**
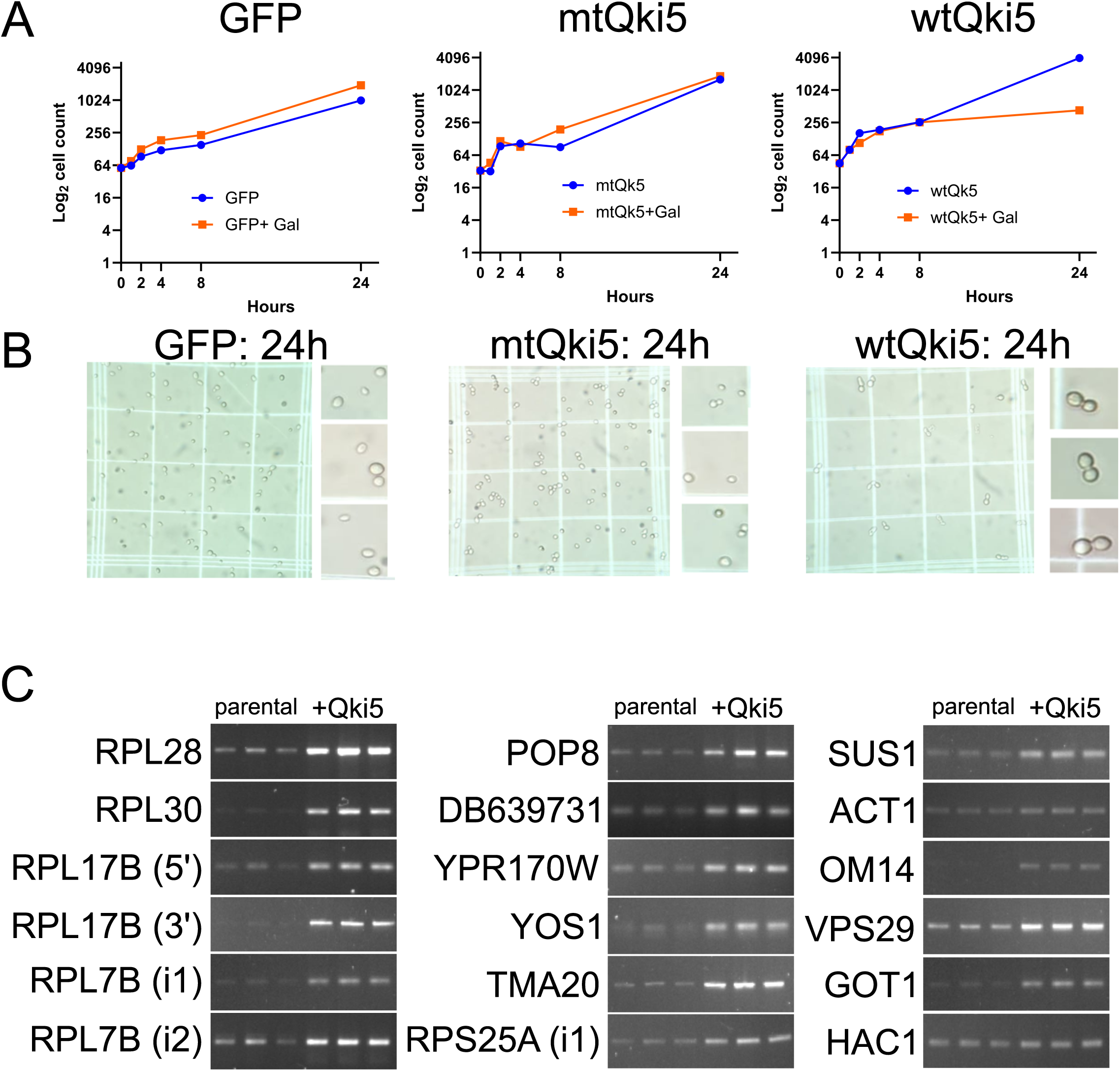
A. Growth curve of Gal-inducible *EGFP*, mt*QKI5* and WT *QKI5* BY4741 yeast cells grown in the absence (blue) or presence of galactose (orange). The y-axis represents the log_2_ number of cells, and the horizontal axis indicates the time point (in hours) at which the cells were collected and counted. B. Representative phase contrast microscopy showing images of EGFP, mtQKI5 and WT QKI5 expressing yeast cells (1000x) at 24 hours after galactose induction. The inset shows enlarged regions to provide more detailed information on cell morphology. C. RT-PCR of parental BY4741 or BY4741 with the *QKI5* transgene 4h after galactose induction, measuring various intron-retention events predicted upon ectopic *QKI5* expression (with exception of control HAC1) using primers that span intron-exon junction for each target and analyzed on an agarose gel (n = 3 per condition).

